# A Reinterpretation of the Relationship Between Persistent and Resurgent Sodium Currents

**DOI:** 10.1101/2023.10.25.564042

**Authors:** Samuel P. Brown, Ryan J. Lawson, Jonathan D. Moreno, Joseph L. Ransdell

## Abstract

The resurgent sodium current (I_NaR_) activates on membrane repolarization, such as during the downstroke of neuronal action potentials. Due to its unique activation properties, I_NaR_ is thought to drive high rates of repetitive neuronal firing. However, I_NaR_ is often studied in combination with the persistent or non-inactivating portion of sodium currents (I_NaP_). We used dynamic clamp to test how I_NaR_ and I_NaP_ individually affect repetitive firing in adult cerebellar Purkinje neurons from male and female mice. We learned I_NaR_ does not scale repetitive firing rates due to its rapid decay at subthreshold voltages, and that subthreshold I_NaP_ is critical in regulating neuronal firing rate. Adjustments to the Nav conductance model used in these studies revealed I_NaP_ and I_NaR_ can be inversely scaled by adjusting occupancy in the slow inactivated kinetic state. Together with additional dynamic clamp experiments, these data suggest the regulation of sodium channel slow inactivation can fine-tune I_NaP_ and Purkinje neuron repetitive firing rates.

**Significance Statement:** Across neuronal cell types, the resurgent sodium current (I_NaR_) is often implicated in driving high rates of repetitive firing. Using dynamic clamp, we determined I_NaR_ is ineffective at driving subsequent action potentials, and that the subthreshold persistent sodium current (I_NaP_) is the critical parameter for scaling repetitive firing rates. We propose I_NaR_ measured in native neurons may reflect a mechanism by which the magnitude of I_NaP_ is fine-tuned.

## Introduction

Purkinje neurons are the exclusive output cells of the cerebellar cortex, delivering GABA-mediated inhibitory signals to neurons in the deep cerebellar nuclei (Ito, 1984; Ito et al., 1964; Obata et al., 1967, 1970). In the absence of synaptic drive, Purkinje neurons have a unique capacity to fire action potentials repetitively at high frequencies (Häusser & Clark, 1997; Raman & Bean, 1997; Thach, 1968), properties which rely on voltage-gated sodium (Nav) channels and the depolarizing currents these channels mediate (Llinás & Sugimori, 1980; Raman & Bean, 1999; Ransdell & Nerbonne, 2018). Beyond the fast inactivating transient sodium current (I_NaT_), Nav channels can generate a persistent sodium current (I_NaP_) component, which was first measured and described in cerebellar Purkinje neurons (Llinás & Sugimori, 1980). Nav channels may also generate a resurgent Nav current (I_NaR_) component, also first characterized in cerebellar Purkinje neurons, which has now been measured in over twenty neuronal cell types throughout the central and peripheral nervous systems (Lewis & Raman, 2014; Raman & Bean, 1997). I_NaR_ is distinct from the transient and persistent Nav currents in that it activates on membrane repolarization and involves the recovery of fast inactivated Nav channels into an open/conducting state (see Figure 1A). In the context of neuronal firing, this type of repolarization would occur during the downstroke of the action potential.

**Figure 1.**
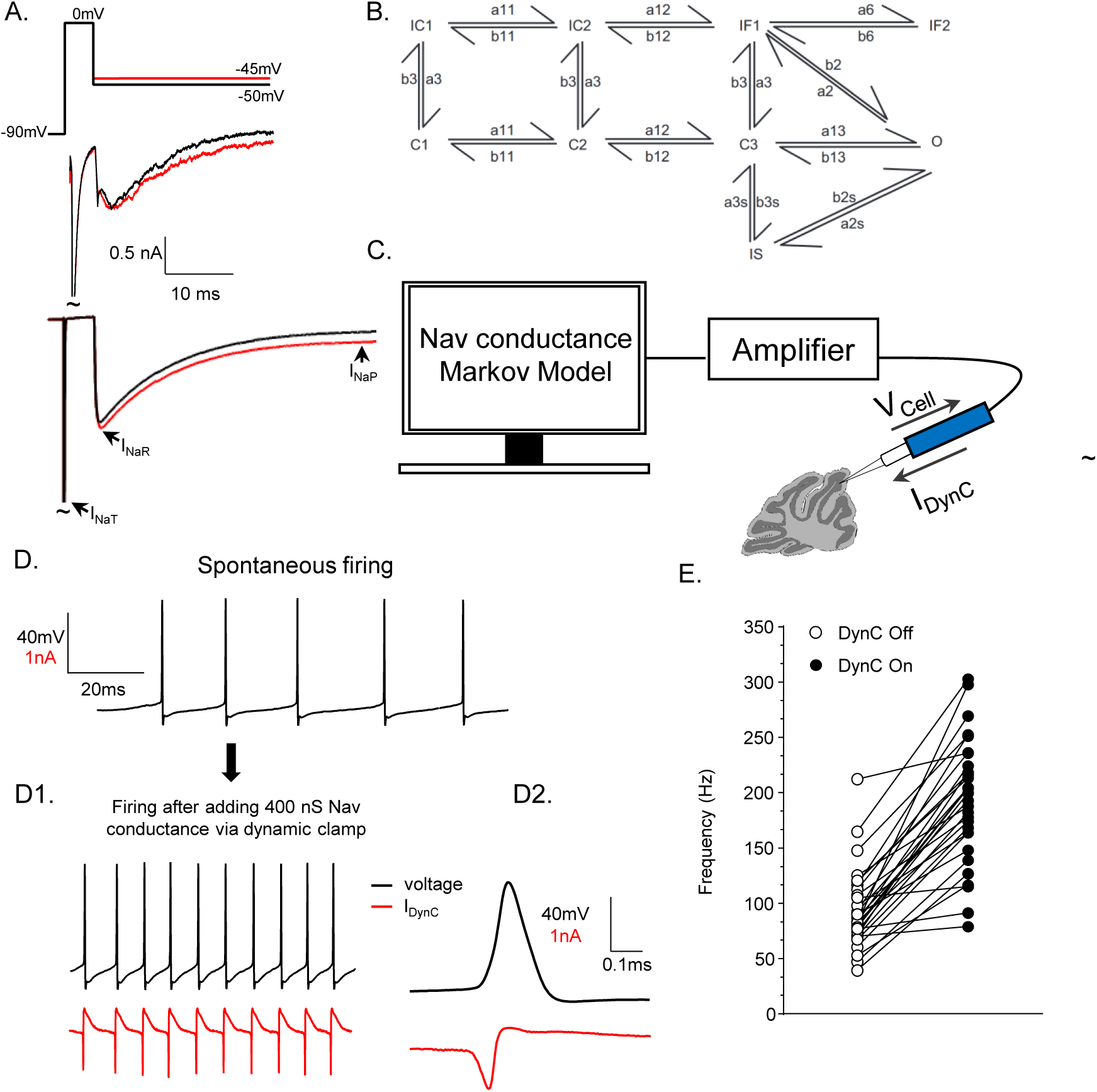
Dynamic clamp addition of a modeled Nav conductance increases the spontaneous firing frequency of adult cerebellar Purkinje neurons **A.** Voltage-clamp step protocol (top) and current traces of the evoked fast-transient (I_NaT_) and resurgent (I_NaR_) Nav currents in an acutely dissociated Purkinje neuron (*middle*). The same voltage protocol was used to evoke and measure Nav currents generated by a Markov Nav conductance model. Labeled arrows depict peak I_NaT_, I_NaR,_ and I_NaP_ on the Nav conductance model trace. **B.** This simulated Nav current was generated by a nine-state gating Markov matrix that includes three closed states (C1, C2, C3), two inactivated-closed states (IC1, IC2), two fast inactivated states (IF1, IF2), a slow inactivated state (IS), and an open state (O). Transition rate constant values are denoted by variables between states (see Methods). Panel **C.** shows a schematic of the dynamic clamp setup used to interface Nav conductance models with a current injection electrode that is patched on an adult Purkinje neuron in a sagittal brain section. A computer running SutterPatch acquisition and dynamic clamp software is interfaced with a Sutter digidata/amplifier, which is connected to the patch electrode. The computer running SutterPatch software uses voltage signals measured in the Purkinje neuron to calculate, in real-time (via the Nav conductance model) current injection values. Current injections representing the model Nav conductance are then applied to the patched cell. **D.** Spontaneous action potentials recorded from an adult Purkinje neuron before and after (**D1**) the dynamic clamp-mediated addition of the Nav conductance represented in panel B. Dynamic clamp-mediated current injections are shown in red. **D2.** Shows an expanded time trace of a single action potential (*black*) and its corresponding dynamic clamp current injection (*red*). **E.** Firing frequency of adult Purkinje neurons (*open circles*) was significantly (P < 0.0001, paired Student’s t-test, n = 30) increased after dynamic clamp-mediated addition of the nominal Nav conductance (*closed circles*). Dynamic clamp-mediated addition of Nav conductances also caused an increase in evoked repetitive firing frequencies (shown in extended data Figure 1-1). Changes in repetitive firing frequency after the addition of the nominal Nav conductance did not correlate with cell passive membrane properties (extended data Figure 1-2).

On its discovery, I_NaR_ was proposed to support high rates of neuronal firing by reducing the Nav channel refractory period (Khaliq et al., 2003; Raman & Bean, 1997, 1999), and this assumption continues to be widely accepted (Quattrocolo et al., 2021; Raman, 2023). During an action potential repolarization, if a subset of Nav channels recover into the open state allowing resurgent Na^+^ influx, the depolarizing current should drive subsequent action potentials. After its discovery, perturbations to I_NaR_ were also associated with changes in repetitive firing (Khaliq et al., 2003; Lewis & Raman, 2014; Raman et al., 1997; Ransdell et al., 2017). For instance, targeted knockout of Navβ4, a sodium channel accessory subunit, was found to result in significantly reduced I_NaR_ in mouse Purkinje neurons (Ransdell et al., 2017) as well as striatal medium spiny neurons (Miyazaki et al., 2014). The loss of Navβ4 was additionally associated with reduced repetitive firing rates compared to wild type controls in these cell types (Miyazaki et al., 2014; Ransdell et al., 2017, 2022). The isolated contribution of I_NaR_ to neuronal repetitive firing, however, has not been directly tested in native neurons.

We use dynamic clamp and multiple (10) Nav conductance models to systematically assess how adjusting properties of I_NaR_ and I_NaP_ affect the firing properties of mouse cerebellar Purkinje neurons. We learned that scaling the proportion and/or kinetic properties of I_NaR_ does not significantly affect Purkinje neuron firing frequency, nor does the addition of I_NaR_ meaningfully impact the action potential waveform in Purkinje neurons. Using voltage-clamp simulation studies, we determined I_NaR_ does not influence repetitive firing rates due to its rapid decay at subthreshold voltages. Subthreshold I_NaP_, however, was found to be a critical parameter in tuning Purkinje neuron repetitive firing rates. In examining the Nav Markov models, we determined peak I_NaR_ and I_NaP_ can be inversely scaled by adjusting occupancy in the slow inactivated kinetic state, an inactivation pathway which is distinct from the conventional fast inactivation pathway. Combined, these data suggest I_NaR_ may reflect a population of Nav channels which would, in the absence of slow inactivation, contribute to I_NaP_; suggesting Nav channel slow inactivation may function to fine tune the amount of Nav channels that contribute to I_NaP_ in order to regulate repetitive firing rates.

## Methods

### Animals

All animal experiments were performed in accordance with protocols approved by Miami University Institutional Animal Care and Use Committee (IACUC) guidelines. All experiments utilized male and female C57BL/6J wild type mice 4-8 weeks old.

### Preparation of acute cerebellar slices

Mice were anaesthetized using an intraperitoneal injection of 1 mL/kg ketamine (10 mg/mL)/xylazine (0.25 mg/mL) cocktail and perfused transcardially with 25 mL cutting solution containing in mM: 240 sucrose, 2.5 KCl, 1.25 NaH_2_PO_4_, 25 NaHCO_3_, 0.5 CaCl_2_, and 7 MgCl_2_. The brain was rapidly dissected, superglued to a specimen tube, and submerged in warmed agarose dissolved in cutting solution. The cerebellum was sliced sagittally in ice-cold cutting solution saturated with 95% O_2_/5% CO_2_ using a Compresstome VF-300 (Precisionary Instruments) at 350 µm thickness. Slices were placed in artificial cerebrospinal fluid (ACSF) containing in mM: 125 NaCl, 2.5 KCl, 1.25 NaH_2_PO_4_, 25 NaHCO_3_, 2 CaCl_2_, 1 MgCl_2_, and 25 dextrose (pH 7.4, ∼300 mOsM) on stretched nylon for 25 minutes at 33°C and then at room temperature for at least 35 minutes before electrophysiological recordings. Purkinje neurons were identified in the Purkinje neuron layer of sagittal cerebellar sections using a SliceScope Pro 3000 (Scientifica).

### Electrophysiological recordings

Whole-cell current-clamp and dynamic clamp recordings were obtained from adult mouse Purkinje neurons (4-8 weeks old) in acutely isolated sagittal cerebellar slices that were constantly perfused with ACSF warmed to ∼34°C and continuously bubbled with 95% O_2_/5% CO_2_. Recording pipettes (boroscilicate standard wall, 1.5 mm outer diameter, 0.86 mm inner diameter) were made using a P-1000 Flaming/Brown Micropipette Puller (Sutter Instruments) and filled with internal solution containing in mM: 0.2 EGTA, 3 MgCl_2_, 10 HEPES, 8 NaCl, 4 Mg-ATP, and 0.5 Na-GTP. Patch pipette resistances were 2-4 MΩ. Prior to patching, pipette tip potentials were zeroed. During current-clamp and dynamic clamp experiments, recorded voltages were corrected for a 17.5 mV liquid junction potential in real-time using SutterPatch software. After a gigaseal was obtained, spontaneous action potentials were recorded using a dPatch Amplifier and SutterPatch software (Sutter Instruments). Input resistance and capacitance calculations were obtained through a whole-cell voltage-clamp protocol in which the membrane voltage was stepped from a -80 mV holding potential to -90 mV (for 100 ms) and to -70 mV (for 100 ms) in a second sweep. This voltage-clamp protocol was applied in each cell prior to current-clamp or dynamic clamp measurements. For membrane capacitance, the integrated area of the capacitive transient current was divided by the change in membrane voltage (10 mV) during each sweep. Action potential properties were analyzed using SutterPatch software (Sutter Instruments). Action potential events were detected if a transient change in membrane potential exceeded 20 mV. Threshold potential was calculated in SutterPatch as the membrane potential prior to an action potential event when the dV/dt is >1 mV/100 µs, or when 25% of the maximum action potential dV/dt is reached, whichever is smaller. Action potential duration was calculated as the time interval from the pass of threshold voltage during action potential upstroke until the pass of threshold voltage during action potential downstroke. Each action potential measurement from a given cell was recorded as the average of the measurement across 20 spontaneous action potentials. Dynamic clamp current injection values, measured during the interspike interval (presented in Figures 5D, 6B), were obtained by measuring the dynamic clamp current injection value 3 ms before the action potential threshold was reached for the subsequent action potential. Action potential peak values were taken at the maximum voltage of the action potential being analyzed. These measurements were made 1 second into each dynamic clamp recording. Recordings were acquired and analyzed using SutterPatch (Sutter Instruments), Microsoft Excel (Microsoft), and Prism (GraphPad) software.

### Markov modeling

A previously developed Markov kinetic state model of Nav channel gating (Ransdell et al., 2022) was used as the base/control model, and is referred to as the ‘nominal model’. To develop the ‘I_NaR_ increased’, ‘I_NaR_ decreased’, ‘τdecay increased’, ‘τdecay decreased’, ‘I_NaR_ reduced’, and ‘I_NaP_ reduced’ models (Figures 2-4), rate constants in the nominal model were altered to cause selective changes in Nav current properties. Each of these models consist of a nine state Markov matrix that includes three closed states (C1, C2, C3), two inactivated-closed states (IC1, IC2), two fast inactivated states (IF1, IF2), a slow inactivated state (IS), and an open state (O). The hand-tuned ‘I_NaR_ increased’ and ‘I_NaR_ decreased’ models, presented in Figure 2, were generated by altering rate constant equations (see Table 1) which resulted in voltage dependent rate constant alterations. For instance, the b3s rate constant changed from 0.94767 to 3.76696 at -45 mV in the ‘I_NaR_ increased’ model and in the ‘I_NaR_ decreased’ model, the b2 rate constant changed from 0.1081 to 3.884 at -45 mV. In order to generate additional models, properties of the originally developed (nominal) model were modified by adjusting values in a parameter optimization algorithm scripted in MATLAB (MathWorks) and previously described in (Moreno et al., 2016; Ransdell et al., 2022). The rate constant values for the nominal and each manipulated model are described in Table 1. To generate the ‘I_NaR_ reduced’ model (Figure 4), state transition rate calculations a3s, b3s, a2s, and b2s were removed from the Q_Matrix.m script prior to parameter optimization. To generate the ‘I_NaP_ reduced’ model (Figure 4), the variable (ResT_line) within the Na_Matrix_DrugFree.m script was changed from 0.18678*V + 22.92 to 0.18678*V + 38.00. Additionally, in line 113 the Err_Late minimization variable was multiplied by 200. To generate the ‘τ_decay_ decreased’ model, the ResT_line variable was changed to 0.18678*V + 38.00. Both Err_ACT and Err_SSA variables in line 113 of the error minimization equation (part of the parameter optimization script) were also multiplied by a value of 10. To generate the ‘τ_decay_ increased’ model, the ResT_line variable was changed to 0.18678*V + 15.00 and the error minimization variables Err_ACT and Err_SSA were multiplied by 90.

**Figure 2.**
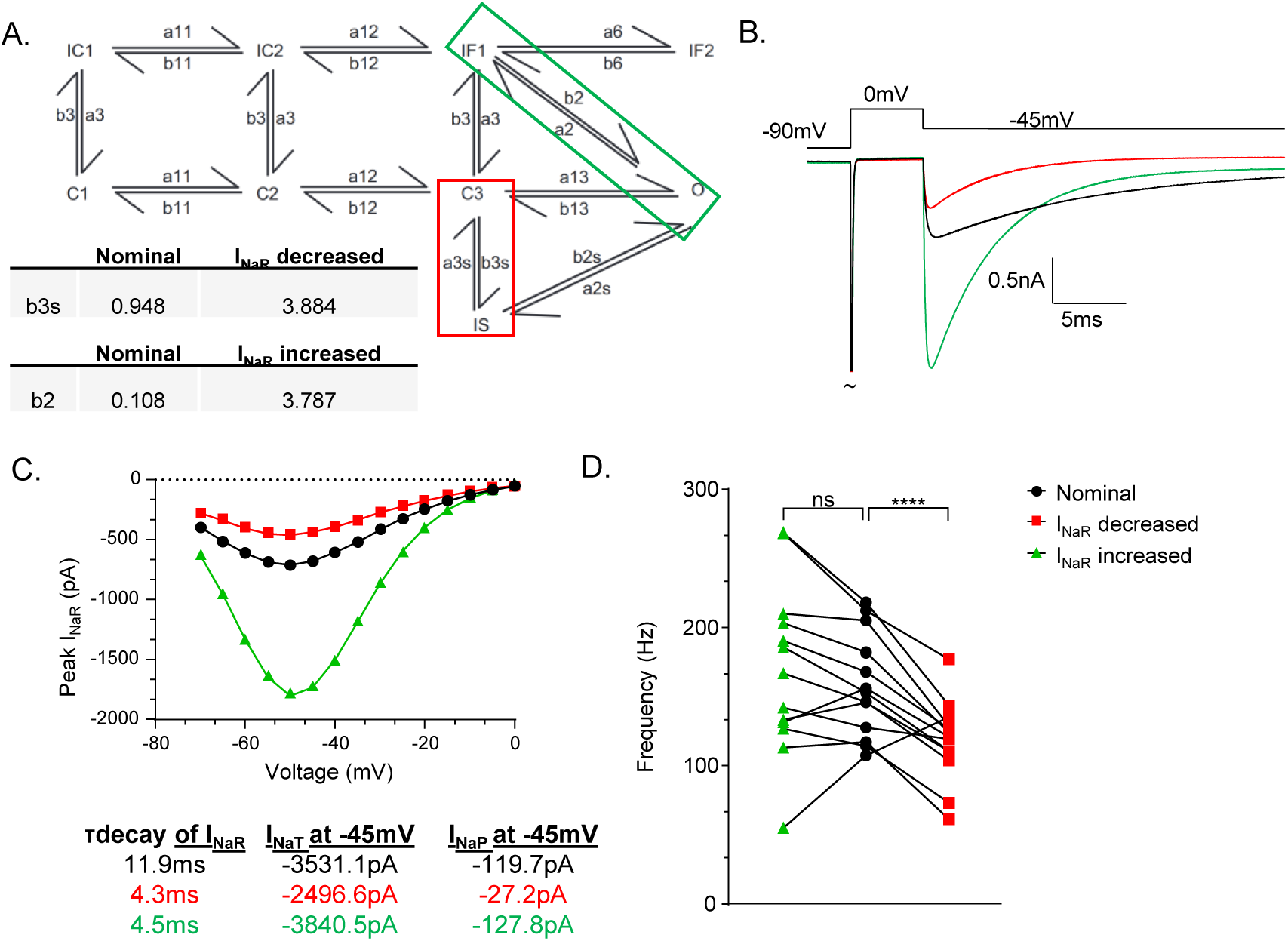
Adjusting peak I_NaR_ does not scale the repetitive firing frequency of Purkinje neurons **A.** To generate the ‘I_NaR_ decreased’ and ‘I_NaR_ increased’ models, transition rate constant values b2 (*green box*) and b3s (*red box*) were selectively adjusted. Values of the nominal and manipulated state transition variables at -45 mV are listed below the matrix. **B.** A simulated voltage-clamp recording of evoked I_NaR_ (at -45 mV) is shown for the nominal model (black), the ‘I_NaR_ increased’ model (*green*), and the ‘I_NaR_ decreased’ model (*red*). **C.** Peak I_NaR_ values were measured at various repolarizing voltage steps after a 5 ms depolarization step to 0 mV. These peak current values are plotted against voltage for the nominal model (black), ‘I_NaR_ increased’ model (*green*), and the ‘I_NaR_ decreased’ model (*red*). **D.** Spontaneous firing frequency of adult Purkinje neurons after dynamic clamp-mediated addition of the nominal (*center, black circles*), ‘I_NaR_ increased’ model (*left, green triangles*), and ‘I_NaR_ decreased’ model (*right, red squares*). Spontaneous firing frequency of the ‘I_NaR_ decreased’ model is significantly reduced compared to spontaneous firing frequencies measured during application of the nominal model. Applying the ‘I_NaR_ increased’ model resulted in no significant change in firing frequency (paired Student’s t-test, ‘I_NaR_ decreased’ vs nominal: P = 0.0042, n = 13; ‘I_NaR_ increased’ vs nominal: P = 0.21, n = 13).

**Figure 3.**
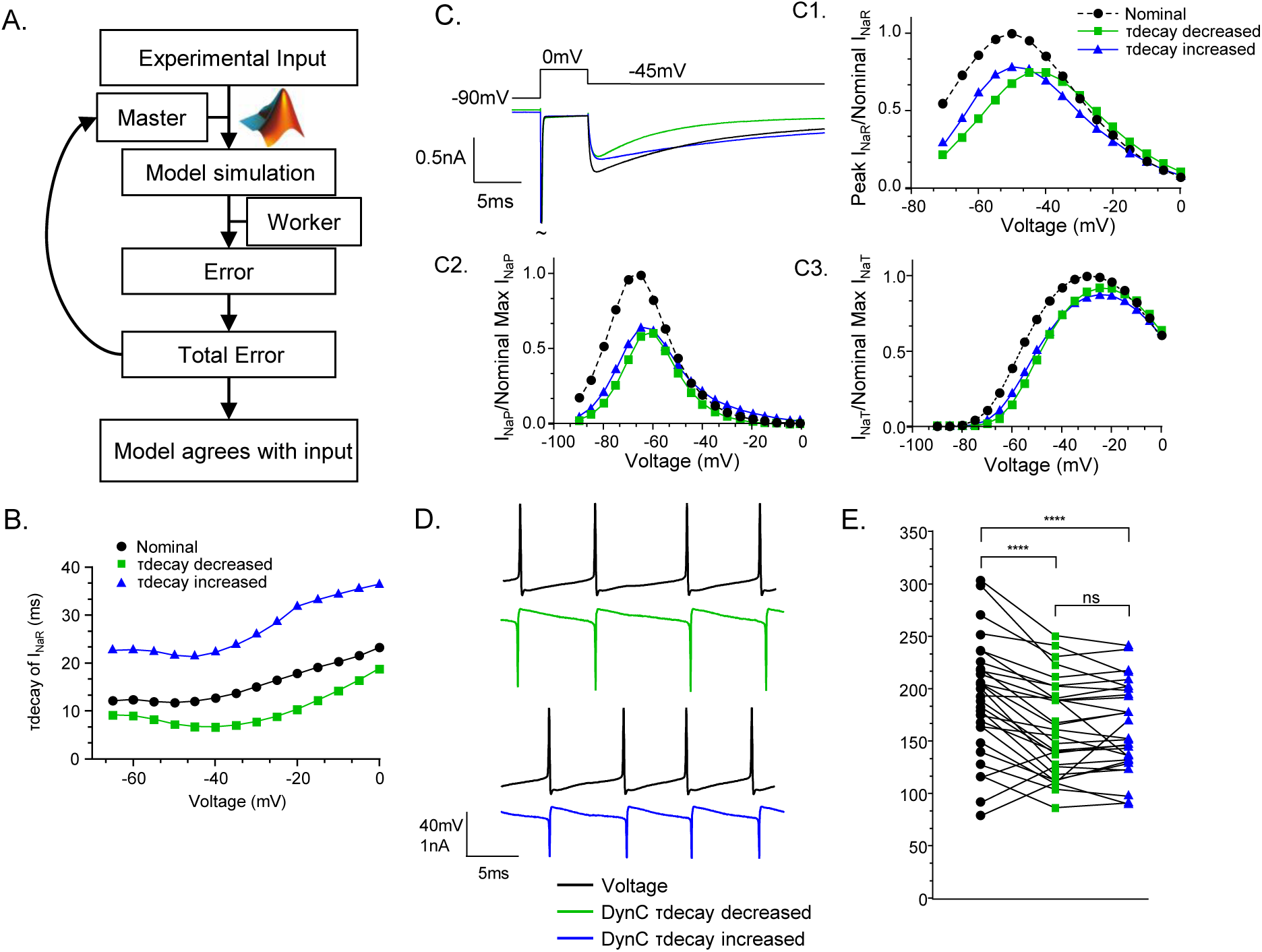
Purkinje neuron repetitive firing rates are not affected by adjusting the time constant of I_NaR_ decay. An automated optimization procedure was used to apply targeted changes to Nav conductance models while preserving other aspects of the nominal Nav conductance model (see Methods). A schematic of this process is shown in panel **A.** A MATLAB script was used to apply the ‘amoeba’ algorithm, checking the difference between the simulation prediction and the target experimental values with each iteration, and ‘crawling’ through potential rate constant values in order to minimize total error. **B.** This amoeba method was used to generate two new Markov conductance models with targeted changes to the time constant of I_NaR_ decay. The time constant of I_NaR_ decay (τdecay) is plotted against voltage for the original nominal model and the two newly generated models, ‘τdecay decreased’ (*green squares*) and ‘τdecay increased’ (*blue triangles*). **C.** A simulated voltage-clamp recording of model generating I_NaR_ is shown for the nominal (*black*), ‘τdecay decreased’ (*green*), and ‘τdecay increased’ (*blue*) models. **C1.** For each model, peak I_NaR_ values are normalized to the max I_NaR_ generated by the nominal model and are plotted against voltage. Similar plots of normalized I_NaP_ and I_NaT_ are presented in panels **C2.** and **C3.**, respectively. **D.** Representative traces showing dynamic clamp-mediated addition of the ‘τdecay reduced’ (*green*) and ‘τdecay increased’ (*blue*) conductances in the same cell. Voltage records are shown in black with the corresponding dynamic clamp current injections shown below. **E.** Spontaneous firing frequencies measured in Purkinje neurons after the addition of the nominal (*black circles*), ‘τdecay decreased’ (*green squares*), and ‘τdecay increased’ (*blue triangles*) model conductances (400 nS) are plotted with lines connecting measurements taken from the same cell. Adding the ‘τdecay decreased’ and the ‘τdecay increased’ model conductances resulted in spontaneous firing frequencies that were significantly reduced compared to firing frequencies measured after adding the nominal model conductance, however, the addition of ‘τdecay decreased’ and the ‘τdecay increased’ model conductances resulted in similar repetitive firing frequencies (paired Student’s t-test-τdecay decreased vs nominal, P < 0.0001, n = 29; τdecay increased vs nominal, P < 0.0001, n = 29; τdecay increased vs τdecay decreased, P = 0.94, n = 29).

**Figure 4.**
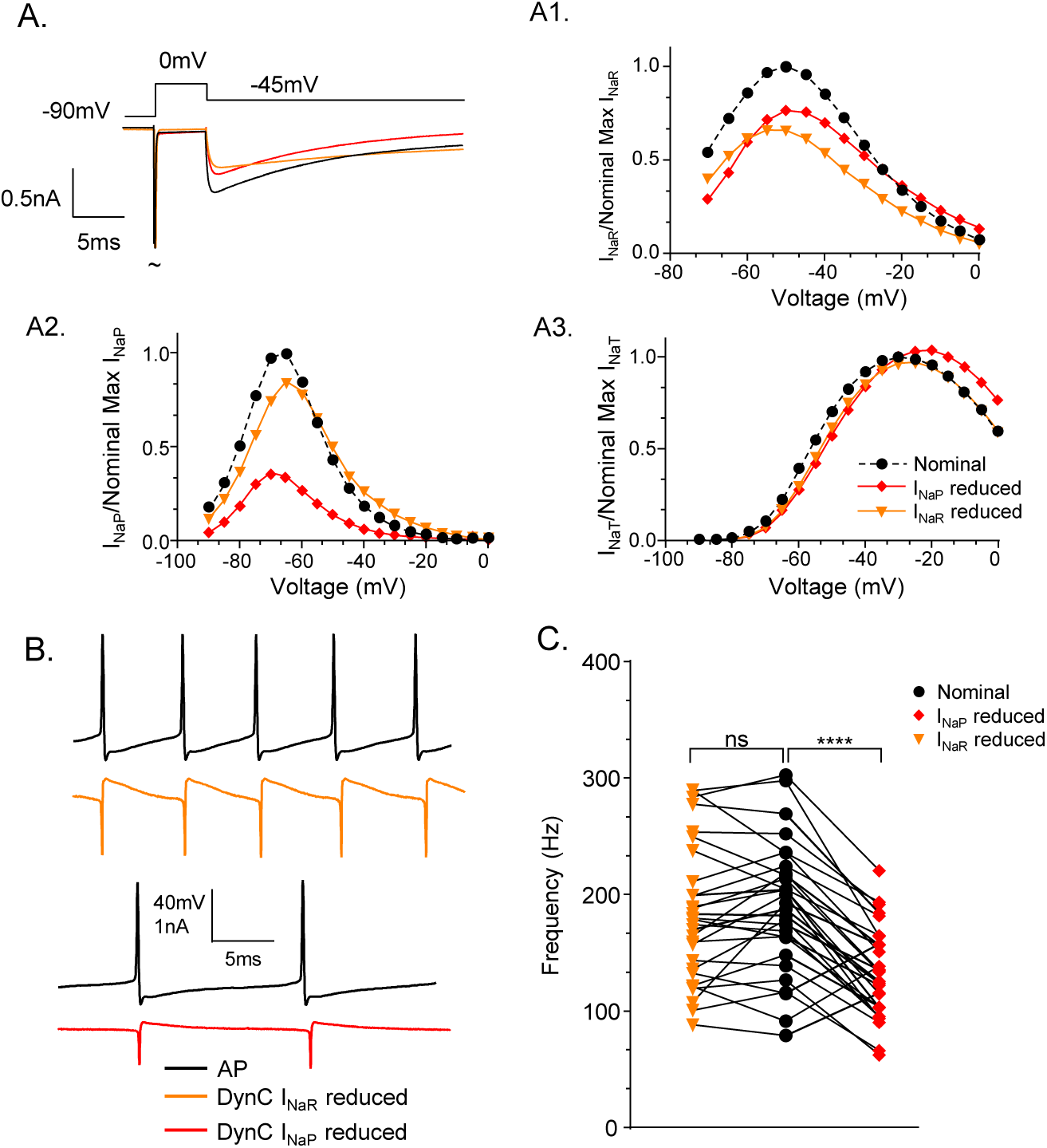
Reducing the amount of I_NaP_ in the modeled conductance significantly decreases spontaneous firing frequency compared to the nominal model, while reducing I_NaR_ does not. **A.** Voltage-clamp simulation traces showing Nav currents generated by the nominal (*black*), ‘I_NaP_ reduced’ (*red*), and ‘I_NaR_ reduced’ (*orange*) models are shown with the common voltage command shown above. **A1-A3.** Peak I_NaR_ (A1), I_NaP_ (A2), and I_NaT_ (A3) were normalized to the peak Nav currents generated by the nominal model and are plotted against voltage. **B.** Representative action potential firing during dynamic clamp-mediated addition of the ‘I_NaR_ reduced’ (*upper*) or ‘I_NaP_ reduced’ (*lower*) conductance models is shown from the same Purkinje neuron. Dynamic clamp current injections are presented as orange or red traces. **C.** The spontaneous firing frequency of Purkinje neurons with addition of the nominal (*black circles*) conductance model was not significantly changed after addition of the ‘I_NaR_ reduced’ conductance model (*left, orange triangles*). However, this firing frequency was significantly decreased after the addition of the ‘I_NaP_ reduced’ conductance model (*right, red diamonds*). (Paired Student’s t-test. nominal vs ‘I_NaR_ reduced’, P = 0.21, n = 29; nominal vs ‘I_NaP_ reduced’, P < 0.0001, n = 29).

**Figure 5.**
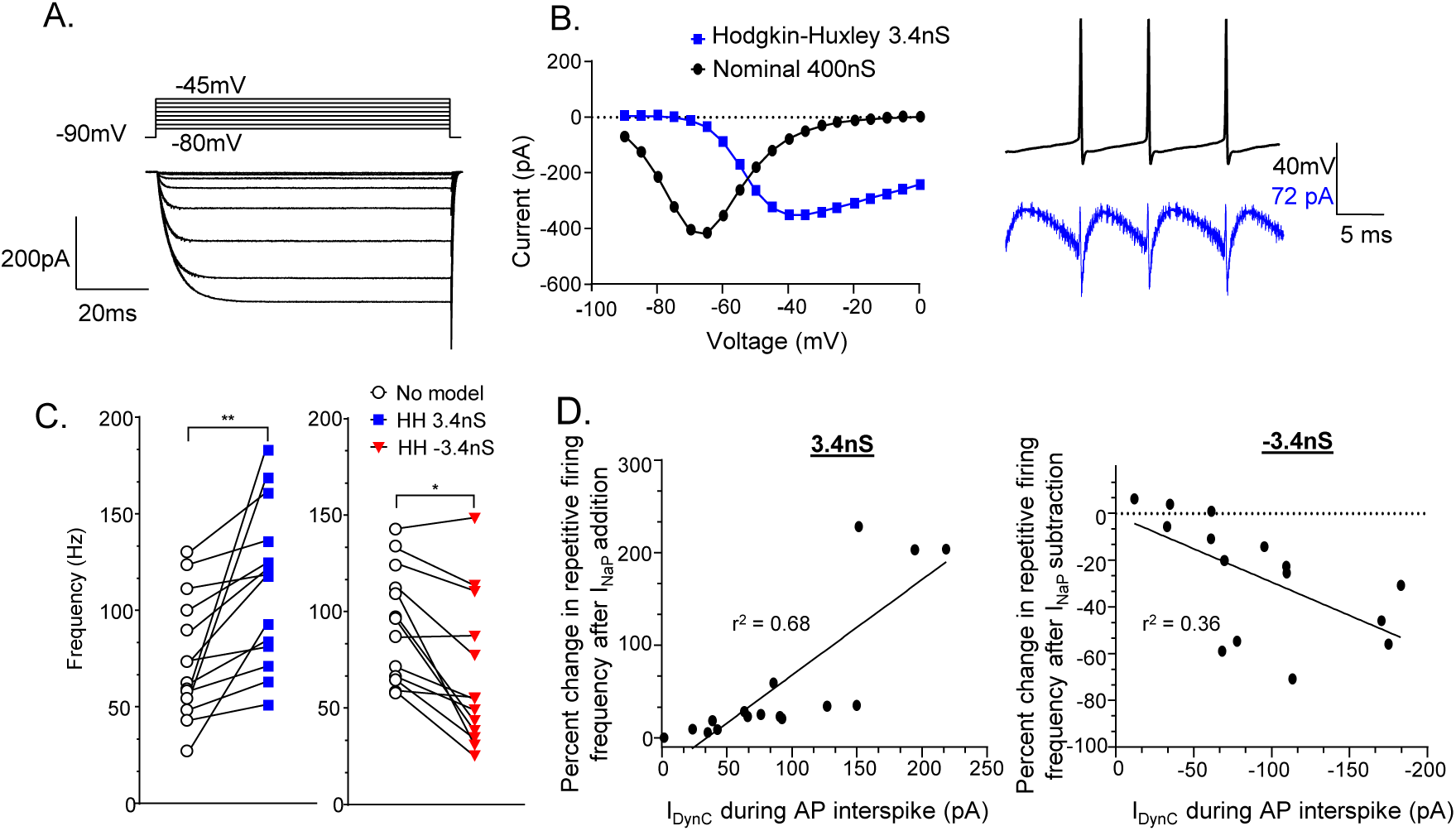
Applying or subtracting an isolated I_NaP_ conductance scales repetitive firing in Purkinje neurons **A.** A Hodgkin-Huxley (HH) modeled conductance (using 3.4 nS conductance) was used to simulate isolated I_NaP_. The current traces presented in panel A. show voltage-clamp of this modeled conductance. **B.** Simulated peak I_NaP_ values versus voltage steps generated by the nominal conductance model (*black*, 400 nS) are compared directly to the HH model (*blue*, 3.4 nS). The HH model’s conductance values were adjusted so that peak I_NaP_ measurements would match the peak I_NaP_ generated by 400 nS of the nominal model. An example of applying the HH model in dynamic clamp experiments is shown to the right; the dynamic clamp current injection is presented in blue. **C.** Spontaneous firing frequency of Purkinje neurons with no dynamic clamp (*open circles*) was significantly increased after dynamic clamp addition of 3.4 nS of the HH conductance model (*left, blue squares*) and significantly decreased after addition of -3.4 nS of the HH conductance model (*right, red triangles*) (Paired Student’s t-test. no model vs 3.4 nS, P = 0.0009, n = 13; no model vs -3.4 nS, P = 0.047, n = 14). **D.** The percent change in repetitive firing frequency after addition (*left*, 3.4 nS) or subtraction (*right*, -3.4 nS) of the HH conductance model is plotted against the amplitude of dynamic clamp current injected during the interspike interval (see Methods) (3.4 nS, r^2^ = 0.68, n = 16; -3.4 nS, r^2^ = 0.36, n = 15).

**Table 1.** Table of rate constants for Markov Nav conductance models.

| Variable | Nominal | $\tau_{\text{decay}}$<br>Reduced | $\tau_{\text{decay}}$<br>Increased | $I_{\text{NaR}}$ Reduced | $I_{\text{NaP}}$ Reduced |
| --- | --- | --- | --- | --- | --- |
| a11 var1 | 0.023989 | 0.02288 | 0.0234922 | 0.024273 | 0.017821 |
| a11 var2 | 961.08 | 962.9323 | 955.288069 | 961.1444 | 955.74551 |
| a12 | 856.13 | 861.1349 | 855.83844 | 855.6402 | 943.691619 |
| a13 | 72.682 | 70.27949 | 75.02708 | 72.05751 | 45.1006 |
| b11 var1 | 0.17233 | 0.13391 | 0.1957689 | 0.145184 | 0.162969 |
| b11 var2 | 19.691 | 18.23159 | 18.08988 | 20.02451 | 17.5219955 |
| b12 | 88.549 | 87.66816 | 90.404 | 88.7198 | 77.35325 |
| b13 | 148.41 | 162.7141 | 230.9623 | 174.5183 | 160.84845 |
| a3 var1 | 0.36734 | 0.514279 | 0.391971 | 0.308533 | 1.006506 |
| a3 var2 | 980.34 | 964.714 | 979.85264 | 980.6689 | 985.713 |
| b3 var1 | 53.241 | 53.5899 | 53.2644 | 53.28371 | 74.161099 |
| b3 var2 | 14.204 | 13.08476 | 13.65378 | 14.39238 | 14.43037 |
| a2 var1 | 87.852 | 84.88214 | 99.3117 | 87.57206 | 83.96448 |
| a2 var2 | 999.72 | 998.5897 | 998.65724 | 999.7462 | 999.9939 |
| a6 var1 | 609.21 | 589.9252 | 621.0904 | 610.1094 | 619.312417 |
| a6 var2 | 86.49 | 84.00415 | 88.18935 | 101.533 | 96.412457 |
| b6 var1 | 156.45 | 224.2441 | 150.19414 | 156.3997 | 147.127477 |
| b6 var2 | 30.317 | 28.8311 | 39.98108 | 58.02996 | 56.26988 |
| a2s var 1 | 15.817 | 37.25738 | 12.52364 | 15.71892 | 22.593552 |
| a2s var 2 | 999.82 | 999.9025 | 999.95304 | 999.7914 | 987.9528 |
| b2s var1 | 0.001001 | 0.001207 | 0.001026 | 0.001003 | 0.001002 |
| b2s va2 | 11.963 | 10.81732 | 13.212289 | 11.96024 | 15.16399 |
| a3s var1 | 0.00447733 | 0.001042 | 0.003335 | 0.005083 | 0.00100109 |
| a3s var2 | 999.93 | 999.2872 | 999.97387 | 999.9429 | 999.97908 |

Transition rates are of the form: $a_{11} = T_{\text{factor}} * 1 / (\text{Input}(1) * \exp(-V / \text{Input}(2)))$ ; $a_{12} = \text{Input}(3) * a_{11}$ ; $a_{13} = \text{Input}(4) * a_{11}$ ; $b_{11} = T_{\text{factor}} * 1 / (\text{Input}(5) * \exp(V / \text{Input}(6)))$ ; $b_{12} = \text{Input}(7) * b_{11}$ ; $b_{13} = \text{Input}(8) * b_{11}$ ; $a_3 = T_{\text{factor}} * \text{Input}(9) * \exp(-V / \text{Input}(10))$ ; $b_3 = T_{\text{factor}} * \text{Input}(11) * \exp(V / \text{Input}(12))$ ; $a_2 = T_{\text{factor}} * (\text{Input}(13) * \exp(V / \text{Input}(14)))$ ; $b_2 = ((a_{13} * a_2 * a_3) / (b_{13} * b_3))$ ; $a_6 = T_{\text{factor}} * (\text{Input}(15) * \exp(V / \text{Input}(16)))$ ; $b_6 = T_{\text{factor}} * \text{Input}(17) * \exp(-V / \text{Input}(18))$ ; $a_{2s} = T_{\text{factor}} * (\text{Input}(19) * \exp(V / \text{Input}(20)))$ ; $b_{2s} = T_{\text{factor}} * (\text{Input}(21) * \exp(-V / \text{Input}(22)))$ ; $a_{3s} = T_{\text{factor}} * \text{Input}(23) * \exp(-V / \text{Input}(24))$ ; $b_{3s} = (a_{2s} * a_{3s} * a_{13}) / (b_{2s} * b_{13})$ ; $Q_{10} = 3$ ; $T = 295$ ; $T_{\text{factor}} = 1.0 / (Q_{10}^{((37.0 - (T - 273)) / 10.0)})$

With the above alterations, each model was generated by running a minimization routine using a Nelder-Mead “amoeba” algorithm-based optimization procedure (Lagarias et al., 1998; Moreno et al., 2016; Nelder & Mead, 1965) to determine the final transition rate constants associated with each manipulated model (presented in Table 1). All models were simulated using a sodium reversal potential of 71.5 mV. All kinetic state occupancy simulations and optimizations were completed in MATLAB 2022a. To verify the Nav currents generated by each model (I_NaT_, I_NaR_, and I_NaP_) had appropriate voltage-dependent and kinetic properties, currents from each model were evoked and measured using steady-state voltage-clamp protocols. Voltage-clamp protocols were applied to an RC circuit (mimicking a passive cell membrane), which was attached to the head-stage while a simulated model conductance was applied via SutterPatch dynamic clamp software. The voltage dependance of activation of I_NaT_ and I_NaP_ were obtained by stepping the model cell (from a -90 mV holding potential) to -70 mV – 0 mV in 5 mV increments for 80 ms. I_NaR_ voltage-dependence of activation was obtained by stepping the model cell from a holding potential of -90 mV to 0 mV for 5 ms, and subsequently applying a repolarizing step to -70 mV to 0 mV (in 5 mV increments) for 80 ms. Prior to analysis of I_NaT_ or I_NaR_, I_NaP_ was digitally subtracted. MATLAB scripts used for the optimization procedure and the nominal Markov state matrix are available on GitHub at https://github.com/morenomdphd/Resurgent_INa. Net charge for Markov model generated currents was measured at steady-state voltage steps by integrating the area under the curve for I_NaT_, I_NaR_, and I_NaP_. I_NaP_ was subtracted prior to net charge calculations of I_NaT_ and I_NaR_. Net I_NaT_ charge was calculated from the beginning of the depolarizing pulse until full decay of the current transient. Net I_NaR_ charge was measured from the beginning of the repolarizing voltage step (after the 5 ms depolarizing prepulse) until 4 * τ (time constant of I_NaR_ decay). Net I_NaP_ charge was also measured during these repolarizing voltage steps across a fixed 10 ms duration, beginning at 4 * τ (time constant of I_NaR_ decay).

### Action potential clamp gating simulations

To visualize gating state probability over time with an action potential as the voltage command, the MATLAB script Fig_4H_4I.m was adjusted (Ransdell et al., 2022). Prior to plotting, the parameter optimization procedure (see previous section) was run to populate the MATLAB workspace with the required variables. An action potential recording, captured from an adult mouse Purkinje neuron firing spontaneous action potentials (at 60.8 Hz) was exported to a Microsoft Excel spreadsheet that contained the time and voltage values of the action potential record. The AP-clamp script was utilized as follows: The script loaded the excel file, which was iterated through the Q_Matrix.m script using the voltage values as the voltage command and the time values as the length of the protocol. Through this script, the values for each gating state as well as I_Na_ and the voltage command were written to the NASIM matrix and then were plotted over time.

### Hodgkin-Huxley modeling

A previously developed Hodgkin-Huxley (HH) model of I_NaP_ in Golgi cells was adopted for dynamic clamp experiments (Solinas et al., 2007) and is available on SenseLab ModelDB (ID: 112685). The conductance value of 3.4 nS was selected for this model because peak HH-generated I_NaP_ values were similar to the peak I_NaP_ from nominal model-generated values at the 400 nS conductance used in other dynamic clamp experiments. The alpha and beta equations for the HH I_NaP_ model are presented below.

| $\alpha$ | $\beta$ |
| --- | --- |
| $5/((0.0421*(V+50)/(1-\exp(-1*(V+50)/5)))+(-0.287*(V+50)/(1-\exp((V+50)/5))))$ | $1/(1+\exp(-1*(V+53)/5))$ |

### Dynamic clamp

Model conductances were applied to whole-cell-patched adult mouse Purkinje neurons through SutterPatch software and a Sutter dPatch amplifier. Markov model conductances were applied in dynamic clamp experiments using a sodium reversal potential of 71.5 mV and a conductance value of 400 nS. The Hodgkin Huxley model, described above, was applied using the same reversal potential and a 3.4 nS conductance. In all dynamic clamp experiments, a model conductance was only applied after a gap-free current-clamp recording (10 second duration) of action potential firing (with dynamic clamp turned off). These intermittent recordings (between applications of model Nav conductances) served as control measurements of Purkinje neuron firing rate without the addition of a model Nav conductance. In all cells included in these analyses, baseline repetitive firing frequencies, with no dynamic clamp current injection, did not vary by more than 10% throughout the duration of the experiment.

## Results

### Dynamic clamp addition of a modeled Nav conductance increases the spontaneous firing frequency of adult cerebellar Purkinje neurons

Purkinje neuron voltage-gated sodium (Nav) currents include fast-transient (I_NaT_), persistent (I_NaP_), and resurgent (I_NaR_) components, which can be isolated and measured using steady state voltage-clamp steps. Voltage-clamp records presented in Figure 1A (*middle traces*) were acquired from an acutely isolated Purkinje neuron (P16) and reflect the I_NaT_ and I_NaR_ components. A brief depolarizing voltage step to 0 mV (from a -90 mV holding potential) activate Nav channels, the majority of which quickly inactivate resulting in the transient Nav current (I_NaT_), and a repolarization voltage step to more negative potentials enable a portion of the fast inactivated Nav channels to recovery into an open/conducting state, which results in ‘resurgent’ sodium influx (I_NaR_). I_NaR_ (at fixed voltages) decays much slower than I_NaT_. At voltages that enable peak I_NaR_ activation (typically around -45 mV), the time constant of I_NaR_ decay is ∼12 ms, and at more hyperpolarized voltages, the time constant of I_NaR_ decay falls to 2-3 ms (Raman & Bean, 1997; Zemel et al., 2021). I_NaR_ decay eventually results in a steady-state depolarizing Nav current in which a population of Nav channels remain in the open/conducting state. This remaining persistent current component reflects I_NaP_. I_NaR_ was first measured in cerebellar Purkinje neurons (Raman & Bean, 1997) and because I_NaR_ is a depolarizing current activated by membrane repolarization, such as the downstroke of an action potential, this depolarizing Nav current is hypothesized to be associated with supporting high rates of repetitive firing. To test this role directly, we used a Markov kinetic state model (depicted in Figure 1B), which accurately reproduces the voltage and time dependent properties of Purkinje neuron Nav currents (Ransdell et al., 2022), to test directly how I_NaR_ properties affect Purkinje neuron repetitive firing. Development of the Nav conductance model is published in Ransdell et al. (2022) and is based on Nav current properties recorded from C57BL/6 mouse Purkinje neurons. The kinetic state Markov model has a single open/conducting state along with two parallel inactivation pathways which are separated as fast (IF1+IF2) and slow (IS) inactivation kinetic states.

To test and verify the dynamic clamp configuration (depicted in Figure 1C), we used SutterPatch dynamic clamp software to apply the model Nav conductance to a passive model cell (Sutter Instruments). Voltage-clamp commands were applied to the model cell with the Markov model conductance added using dynamic clamp software, and the model-generated currents were recorded. Nav currents generated by this conductance model have similar voltage-dependent and kinetic properties as those measured in mouse Purkinje neurons (Ransdell et al., 2022). Figure 1A (bottom) current traces show the performance of the Nav conductance model under an identical voltage command. The nominal conductance model produces I_NaT,_ I_NaR_, and I_NaP_ current components. We utilized this model throughout our dynamic clamp experiments as a baseline addition of Nav conductance and refer to it as the ‘nominal’ model throughout the rest of this manuscript. The nominal model conductance was applied to adult (4-7-week-old) Purkinje neurons in acute sagittal brain slices at physiological temperatures. The nominal model applied at 400 nS (see Methods) significantly (P < 0.0001, paired Student’s t-test, n = 30) increased the spontaneous firing frequency of Purkinje neurons, compared to spontaneous firing without dynamic clamp (Figure 1E). Addition of the nominal model conductance also resulted in significant (P < 0.001, RM one-way ANOVA, n = 24) increases in evoked firing frequency during depolarizing current injections (see extended data Figure 1-1).

### Adjusting peak I_NaR_ does not scale the repetitive firing frequency of Purkinje neurons

We developed and tested Nav conductance models that have varying levels of peak I_NaR_. To alter peak I_NaR_, individual rate constants in the nominal Markov model (presented in Figure 1) were adjusted. By increasing the rate constant b2, which is responsible for transiting channel occupancy from the fast inactivated state into the open state, peak I_NaR_ was increased to a peak value of 1.38 nA (measured at -45 mV), compared to 680 pA in the nominal model (see Figure 2A, B, *green*). Alternatively, by increasing the b3s rate constant, which is responsible for transiting channel occupancy from the closed state into the slow inactivated state, peak I_NaR_ was reduced to 350 pA (measured at -45 mV) (Figure 2A, B, *red*). Voltage-clamp records of Nav currents produced by these models, along with peak I_NaR_ I-V plots for these models are shown in Figures 2B, and C. To test how scaling I_NaR_, using these new models, affects the repetitive firing properties of Purkinje neurons, we added the model variants to Purkinje neurons using dynamic clamp. The nominal model, along with the ‘I_NaR_ decreased’ (*red*) and ‘I_NaR_ increased’ (*green*) models were applied to the same Purkinje neurons using 400 nS conductance for each model. Consistent with the hypothesis that I_NaR_ supports high rates of repetitive firing, we found adding the ‘I_NaR_ decreased’ model conductance resulted in significantly reduced repetitive firing frequencies compared to addition of the nominal model conductance (Figure 2D, *left*). However, when the ‘I_NaR_ increased’ conductance model was added, there was no significant change in the repetitive firing rates compared to addition of the nominal model conductance (Figure 2D, *right*). The ‘I_NaR_ increased’ conductance model has a peak I_NaR_ value that is 2.3 times larger than the nominal model (see Figure 2C). We reasoned that additional Nav current properties, other than peak I_NaR_, were likely altered in creating the ‘I_NaR_ increased’ and ‘I_NaR_ decreased’ models, which may have influenced the effects of each model conductance on repetitive firing frequencies. Analysis of the voltage-clamp simulation traces revealed the time constant of I_NaR_ decay was reduced in both the ‘I_NaR_ decreased’ and ‘I_NaR_ increased’ models, compared to the nominal model. Additionally, we noted I_NaP_ was much smaller in the ‘I_NaR_ reduced’ model compared to the nominal model. Measures of these Nav current components are presented under the I-V plot in Figure 2C.

### Purkinje neuron repetitive firing rates are not affected by adjusting the time constant of I_NaR_ decay

Because altering a single rate constant was found to alter multiple Nav conductance parameters, notably the tau of I_NaR_ decay and the proportion of I_NaP_, we utilized a numerical optimization procedure (see schematic in Fig. 3A and description in Methods) to develop additional Nav conductance Markov models that exhibit more targeted changes to the nominal model properties, while working to conserve other Nav conductance parameters in the nominal model. Our first goal was to test if the rate of I_NaR_ decay influences how a Nav conductance affects repetitive firing. To develop models with either a longer or shorter I_NaR_ decay, compared to the nominal model, we adjusted the constraint for the decay of I_NaR_ before applying the rate constant optimization (amoeba method) procedure (Figure 3A). This strategy allowed us to develop two new conductance models that, compared to the nominal model in Figure 3B-C, have a tau of I_NaR_ decay that is either longer (‘τdecay increased’, *blue*) or shorter (‘τdecay decreased’, *green*). The time constant of I_NaR_ decay is plotted against voltage for each model in Figure 3B. While these models have different peak I_NaT_, I_NaP_, and I_NaR_ values (using a 400 nS conductance) than the nominal model (see Figures 3C1-C3), the values of these current components were similar to one another. This enabled us to test the effect of altered I_NaR_ decay rates directly using dynamic clamp experiments.

When the two newly developed models, ‘τdecay increased’ and ‘τdecay decreased’ were added at 400 nS to adult Purkinje neurons, both models were found to cause significant increases in repetitive firing, compared to no model. However, the increased repetitive firing rates (under both models) were not significantly different from one another (Figure 3D, E), indicating that while adding the modeled Nav conductance increases repetitive firing rates, applying substantial changes to the tau of I_NaR_ decay does not affect these firing rates (see Figure 3D). Both of the new models resulted in significantly lower firing rates than the nominal model (Figure 3E), however, we also noted that both the ‘τdecay increased’ and ‘τdecay decreased’ models generate lower peak I_NaP_ values than the nominal model (Figure 3C2).

### Peak I_NaP_, not peak I_NaR_, is critical in scaling Purkinje neuron repetitive firing rates

We hypothesized that I_NaP_ may be the critical parameter for scaling Purkinje neuron repetitive firing rates. To test this hypothesis, we again used the constrained numerical optimization method, (presented in Figure 3A) to develop a Nav conductance with reduced I_NaP_. This model is referred to as the ‘I_NaP_ reduced’ model. Even utilizing numerical optimization methods, it was difficult to develop the ‘I_NaP_ reduced’ model without also significantly reducing peak I_NaR_ values (found in the nominal model, see Figure 4A1, *red* and *black*). For this reason, we developed an additional Nav model which has similar levels of I_NaP_ and I_NaT_ as the nominal model (Figure 4A2, 4A3, *orange*), but that also has a reduced peak I_NaR_ compared to the nominal model (Figure 4A1, *orange*). We refer to this model as ‘I_NaR_ reduced’. Dynamic clamp-mediated addition of these models could then be used to compare the effects of reduced I_NaP_ or reduced I_NaR_ on Purkinje neuron firing. Results from these dynamic clamp studies, presented in Figure 4B, C, reveal addition of the ‘I_NaR_ reduced’ model has no effect on repetitive firing rate compared to the nominal model (Figure 4C, *left*) and addition of the ‘I_NaP_ reduced’ model results in significantly (P < 0.0001) lower repetitive firing rates than addition of the nominal model (Figure 4C, *right*).

### Applying or subtracting I_NaP_ conductance alone scales repetitive firing

The nominal model, and subsequent iterations of the Nav conductance models developed in Figures 3 and 4, have a persistent component that activates maximally at around -60 mV (see Figures 3C2, 4A2). However, I_NaP_ measured in Purkinje neurons typically activates at more depolarized voltages, with peak persistent currents evoked at around -40 mV (Ransdell et al., 2017). To test how this shift in the voltage-dependence of I_NaP_ activation affects the contribution of I_NaP_ to repetitive firing in Purkinje neurons, and to also examine how sole application of I_NaP_ affects repetitive firing in Purkinje neurons, we utilized an I_NaP_ Hodgkin-Huxley (HH) model (previously published in Solinas et al., 2007) for dynamic clamp experiments. We tested the I_NaP_ HH model using voltage-clamp simulations and adjusted the conductance of the model to match the peak I_NaP_ values measured from the nominal model (see Figure 5A, B). The HH I_NaP_ conductance model applied in dynamic clamp using 3.4 nS conductance was found to be sufficient to increase Purkinje neuron repetitive firing frequency (Figure 5C, *left*). Additionally, we tested if subtracting this conductance via dynamic clamp reduces repetitive firing. We found that dynamic clamp-mediated subtraction of I_NaP_ results in a significant reduction in repetitive firing frequency (Figure 5C, *right*). As is evident in Figure 5C, the HH I_NaP_ conductance, applied across cells at either 3.4 nS or -3.4 nS, has variable effects on firing frequency. In some cells, adding 3.4 nS I_NaP_ caused only slight increases in firing frequency, while in other cells, adding this conductance drove a nearly 3-fold increase in firing frequency.

We investigated if passive membrane properties might influence the contribution of I_NaP_ to firing frequency. No correlation was found between the input resistance of the cell or capacitance with the increase or decrease in firing frequency after applying 3.4 nS or -3.4 nS, respectively, of the I_NaP_ conductance model (data not shown). These passive membrane properties also did not correlate with the change in cell firing after applying the nominal model conductance (see extended data Figure 1-2). However, dynamic clamp current injection values during the interspike interval, measured 3 ms prior to action potential threshold (see Methods), were found to correlate with the percent increase or decrease in firing frequency after applying the 3.4 nS I_NaP_ model (r^2^ = 0.68) or -3.4 nS I_NaP_ model (r^2^ = 0.36), respectively. These data suggest the steady-state depolarizing current provided by I_NaP_ during interspike intervals is a critical determinant in scaling the repetitive firing rates of Purkinje neurons.

### When applied using dynamic clamp, peak Nav current components (I_NaT_, I_NaR_, and I_NaP_) do not correlate with changes in the action potential waveform

Application of the nominal model to spontaneously firing Purkinje neurons resulted in significant alterations to the action potential duration (increased), peak (increased), amplitude (increased), and threshold voltage (decreased) (see Table 2). The amplitude of the action potential after-hyperpolarization and the maximum dV/dt, taken from a first derivative phase plot of the action potential waveform, were not significantly affected by applying the nominal model conductance (Table 2). Other Markov conductance models had varying effects on these action potential properties. To test if the magnitude of the Nav current components (I_NaT_, I_NaR_, and I_NaP_), measured across models, scales or consistently alters a given action potential property, we plotted the current value of I_NaT_, I_NaR_, and I_NaP_ from each Markov model against the mean change in the action potential parameter after addition of the respective model. These plots are shown in the extended data Figure supporting Table 2 (labeled as Table 2-1), and indicate the size of these Nav current components across models do not consistently affect or scale any of the measured parameters of the action potential waveform.

**Table 2.** Action potential waveform properties after adding each Markov Nav conductance.

| Model | Mean AP duration (100%)<br>(μs) | Mean AHP (mV) | Mean AP Peak (mV) | Mean AP Threshold (mV) | Mean AP Amplitude<br>(mV) | Max dV/dt (V/μs) |
| --- | --- | --- | --- | --- | --- | --- |
| No model (n = 29) | 411.8 ± 11.95 | -73.10 ± 0.561 | 1.024 ± 1.089 | -59.11 ± 0.546 | 60.14 ± 1.242 | 587.7 ± 20.35 |
| Nominal (n = 29) | 475.3 ± 13.20<br>(p < 0.0001) | -72.70 ± 0.615<br>(p = 0.4499) | 5.662 ± 1.044<br>(p < 0.0001) | -56.86 ± 0.578<br>(p = 0.0026) | 62.52 ± 1.329<br>(p = 0.0011) | 582.2 ± 25.62<br>(p = 0.7291) |
| I<sub>NaP</sub> Reduced (n = 37) | 488.6 ± 11.82<br>(p = 0.0015) | -73.41 ± 0.588<br>(p = 0.9242) | 5.544 ± 1.342<br>(p < 0.0001) | -57.08 ± 0.628<br>(p < 0.0001) | 62.62 ± 1.720<br>(p < 0.0001) | 573.7 ± 26.18<br>(p = 0.0001) |
| I<sub>NaR</sub> Reduced (n = 35) | 523.9 ± 33.22<br>(p = 0.0153) | -72.93 ± 0.755<br>(p = 0.6177) | 5.769 ± 1.213<br>(p = 0.8196) | -56.60 ± 0.807<br>(p = 0.0022) | 62.37 ± 1.513<br>(p = 0.6351) | 555.3 ± 24.62<br>(p = 0.0282) |
| τ<sub>decay</sub> decreased (n = 34)<br>Figure 3 | 513.4 ± 17.43<br>(p = 0.0009) | -74.34 ± 0.501<br>(p < 0.0001) | 10.93 ± 1.245<br>(p < 0.0001) | -58.84 ± 0.463<br>(p = 0.1665) | 69.78 ± 1.410<br>(p = 0.0003) | 670.6 ± 24.89<br>(p < 0.0001) |
| τ<sub>decay</sub> increased (n = 36)<br>Figure 3 | 497.9 ± 13.32<br>(p < 0.0001) | -73.48 ± 0.700<br>(p = 0.0601) | 7.036 ± 1.320<br>(p < 0.0001) | -56.98 ± 0.717<br>(p = 0.0318) | 64.02 ± 1.484<br>(p < 0.0001) | 589.5 ± 24.63<br>(p < 0.0001) |
| I<sub>NaR</sub> decreased (n = 17)<br>Figure 2 | 488.3 ± 28.81<br>(p = 0.0183) | -72.50 ± 1.321<br>(p = 0.6783) | 1.364 ± 1.046<br>(p = 0.9883) | -56.29 ± 1.812<br>(p = 0.2675) | 57.65 ± 2.402<br>(p = 0.5275) | 502.7 ± 33.32<br>(p = 0.0968) |
| I<sub>NaR</sub> increased (n = 15)<br>Figure 2 | 508.9 ± 50.16<br>(p = 0.2171) | -72.14 ± 1.498<br>(p = 0.3844) | 3.745 ± 1.112<br>(p = 0.0281) | -52.79 ± 1.872<br>(p = 0.0088) | 56.53 ± 1.817<br>(p = 0.2323) | 478.4 ± 20.75<br>(p = 0.0005) |

Mean ± standard deviation is presented across models for each action potential (AP) waveform parameter. P values are the result of paired Student's t-tests which compare action potential data with no dynamic clamp and action potential data after adding the Markov conductance model (see Methods). AHP refers to action potential after hyperpolarization (see Methods).

### Across dynamic clamp Nav conductance models, I_NaP_ is positively correlated with changes in repetitive firing frequency

Using a similar approach as Figure 5D, we examined how dynamic clamp current injection values relate to changes in repetitive firing frequency across Markov Nav conductance models. We reasoned that during application of a model conductance, the contribution of I_NaP_ is most localized during the interspike interval immediately preceding the subsequent action potential. For each of the seven Markov conductance models, I_DynC_ was measured during the interspike interval (3 ms prior to the threshold potential of the subsequent action potential), and the mean I_DynC_ was calculated for each model. Across the seven Markov models, these mean I_DynC_ values were plotted against the mean percent increase in repetitive firing (Figure 6B) for each model. This plot revealed a strong positive correlation (r^2^ = 0.93), suggesting the steady-state Nav current, which occurs at subthreshold voltages, is a critical parameter for Nav conductance mediated increases in repetitive firing frequency. In contrast, if the I_DynC_ value was measured at the peak of the action potential waveform (see Figure 6A schematic), there was no correlation (r^2^ = 0.004) with each model’s effect on repetitive firing frequency (Figure 6C).

**Figure 6.**
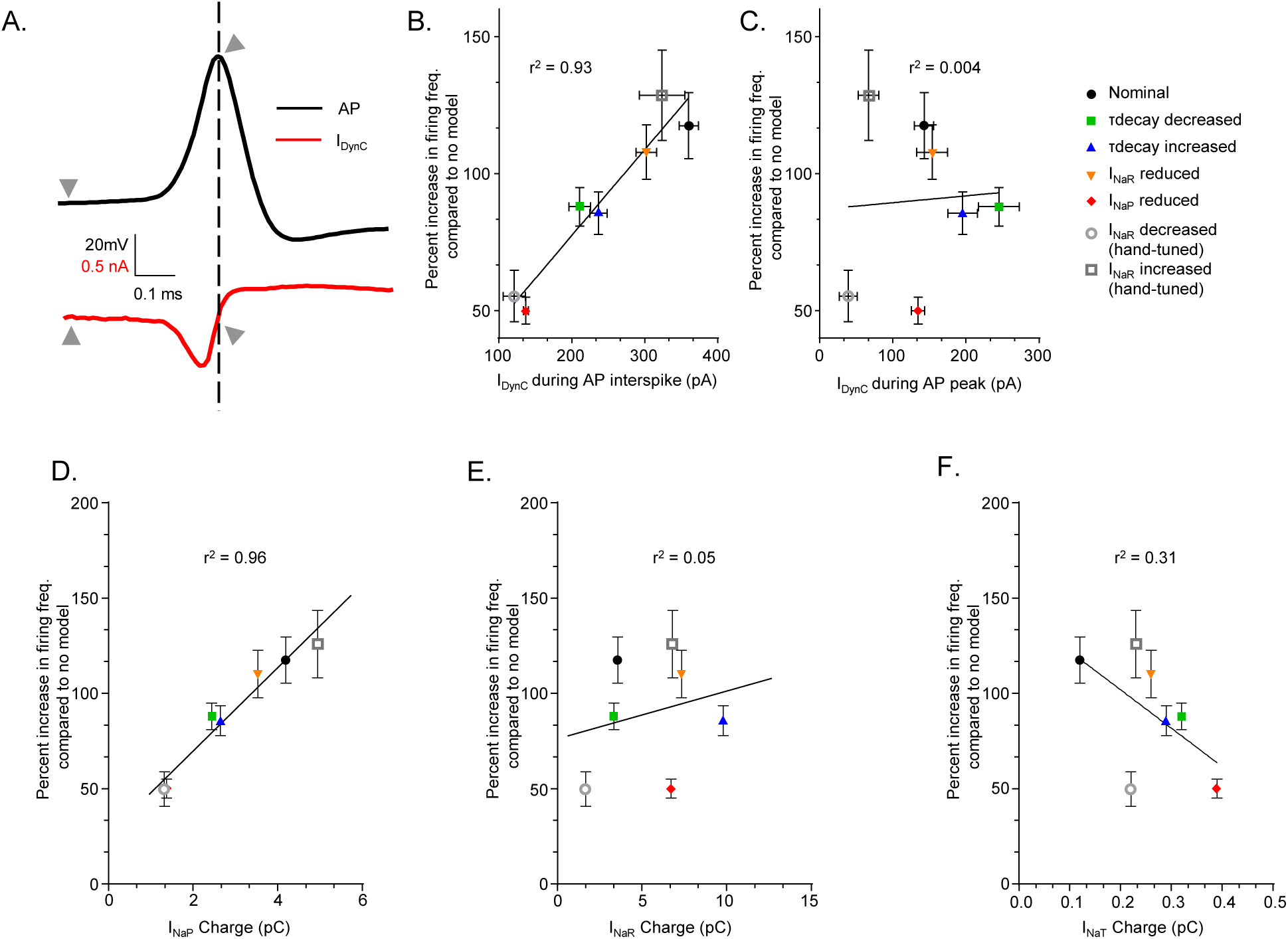
Across models used for dynamic clamp, I_NaP_ is positively correlated with the changes in repetitive firing frequency. The action potential waveform shown in panel **A.** depicts how dynamic clamp current injection values were obtained during the interspike intervals and action potential peaks (see Methods). **B.** A strong correlation exists between the percent increase in spontaneous firing frequency, driven by the addition of each Markov conductance model, and the dynamic clamp current injection during the interspike interval of an action potential (r^2^ = 0.93). If the dynamic clamp current injection is measured at the peak of the action potential, the correlation coefficient (r^2^) falls to 0.004 (**C.**). **D.** There is a strong correlation between the percent increase in spontaneous firing frequency after addition of each Markov conductance model and the net I_NaP_ charge (for each model) during a -65 mV steady-state voltage-clamp step (r^2^ = .96; P = 0.0001). This positive correlation is absent if measurements of net I_NaR_ charge (measured at -45 mV) (**E.**) or net I_NaT_ charge (measured at 0 mV) (**F.**) are used (r^2^ = 0.05, P = 0.61; r^2^ = 0.31, P = 0.19, respectively). Pearson’s correlation tests were used to determine significance. Net charge calculations for each current component are described in Methods.

To examine more directly how each current component affected firing, we plotted for each model the net charge calculated from I_NaT_, I_NaR_, and I_NaP_ records. Net charge (for each Nav current component) was calculated from the current evoked by a single steady-state voltage-step (see Methods). These net charge values were plotted against the mean percent increase in repetitive firing frequency after the model was applied in dynamic clamp. These plots again reveal a strong correlation between I_NaP_ measured at -65 mV and the change in repetitive firing rates (Figure 6D), while I_NaR_ (Figure 6E), and I_NaT_ (Figure 6F) values were not significantly correlated to the mean increases in repetitive firing rate. Similar plots, which compared the mean percent change in repetitive firing rates with the measured peak I_NaP_, I_NaR_, and I_NaT_, values for each model resulted in a similar strong correlation with I_NaP_ (r^2^ = .96) and weak correlations with I_NaR_ and I_NaT_ (r^2^ =.41, and .38, respectively, data not shown).

### Markov model open state occupancy reveals steady-state I_NaP_ is the primary depolarizing drive of subsequent action potentials

Across models, we hoped to determine why scaling peak I_NaR_ does not affect the repetitive firing properties of Purkinje neurons. To analyze this more directly, we used voltage-clamp simulation experiments in which the voltage command consisted of two consecutive action potential waveforms previously recorded from a Purkinje neuron firing at ∼60 Hz. Under this voltage-clamp paradigm, we analyzed the performance of the nominal model in addition to the ‘I_NaR_ reduced’ model (orange), and the ‘I_NaP_ reduced model’ (red) presented in Figure 4. In addition to examining the current evoked by this voltage command for each model, we measured and plotted occupancy in the open/conducting kinetic state. Time-locked traces from this examination are presented in Figure 7. In comparing the ‘I_NaR_ reduced’ model (*orange*) traces with the nominal (*black*) traces, there was only a slight difference in the open-state occupancy (middle traces) and evoked steady-state I_Na_ traces during the interspike interval. This is consistent with the ‘I_NaR_ reduced’ model having a slightly smaller I_NaP_ component than the nominal model (see Figure 4A2). In the ‘I_NaR_ reduced’ model’s current trace and open-state occupancy traces (*orange traces*), we found no evidence of the reduced I_NaR_ component. This is likely because I_NaR_ is very small at subthreshold voltages and because the decay of I_NaR_ is relatively rapid at hyperpolarized membrane potentials. However, the ‘I_NaP_ reduced’ model (*red traces*) has significantly less open-state-occupancy corresponding to less steady-state evoked I_Na_ during the interspike interval, revealing why the ‘I_NaP_ reduced’ model, when applied using dynamic clamp, results in significantly lower firing frequencies than the nominal model and the ‘I_NaR_ reduced’ model (see Figure 4C).

**Figure 7.**
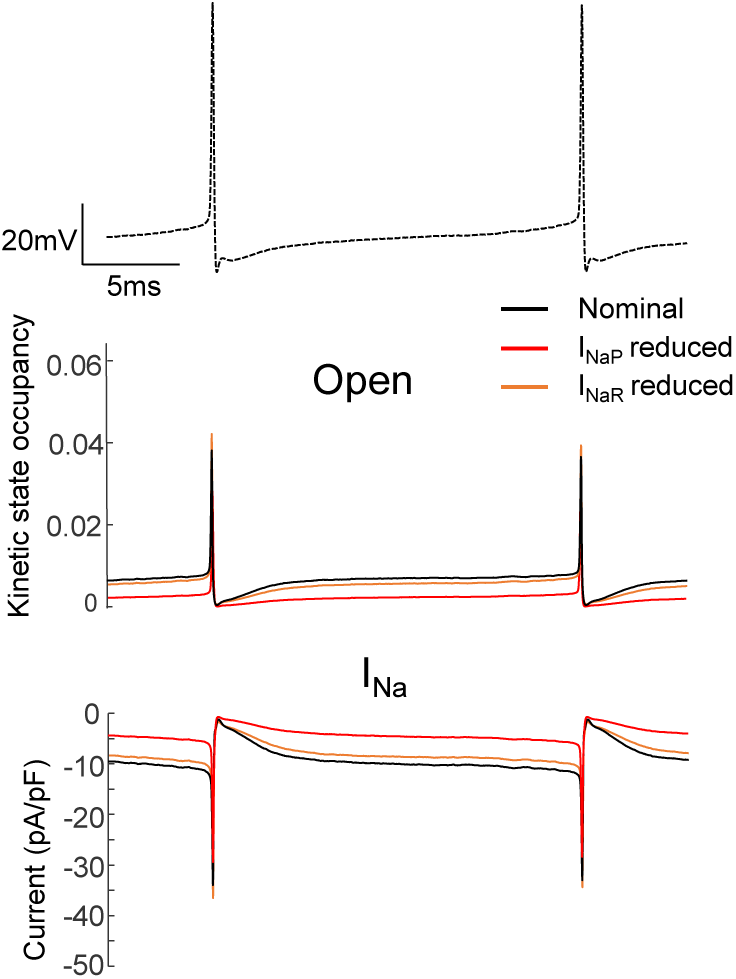
Simulation action potential clamp experiments were conducted using a voltage command (dashed trace, *upper*) that includes two action potentials that were originally recorded from a Purkinje neuron firing at ∼60 Hz. The nominal (*shown in black*), ‘I_NaR_ reduced’ (*shown in orange*), and ‘I_NaP_ reduced’ (*shown in red*) Markov conductance models (see Figure 4) were applied to a model cell in MATLAB (see Methods). Kinetic state occupancy plots, showing occupancy in the open state, are shown below (*middle traces*) and are time-locked with the voltage command. The bottom traces show the time-locked evoked currents produced by the voltage command for each of these models.

### Peak I_NaP_ and I_NaR_ can be inversely adjusted by adjusting occupancy in the slow inactivated kinetic state

In developing the various models with alterations in I_NaP_ and I_NaR_, it became evident that increased occupancy in the slow inactivated kinetic state, shown as IS in the Markov kinetic state models (Figure 8A), was necessary to increase peak I_NaR_. On reflection, this is unsurprising because the decay of the I_NaR_ current is simulated in these Markov models by transiting open channels into the IS state. Any simulated channels that fail to transit from the open state into IS, which is absorbing (Ransdell et al., 2022), are available to contribute to I_NaP_. According to these model properties, we reasoned that we should be able to inversely scale peak I_NaP_ and I_NaR_ by adjusting the rate constants that affect Open and IS state occupancy. These rate constants are labeled b2s and a2s and are highlighted in panel 8A. We conducted voltage-clamp simulation studies to test this hypothesis and indeed found that reducing the b2s rate constant results in an increase in the proportion of I_NaR_ and reduces I_NaP_ (Figure 8B). Alternatively, if the a2s rate constant is reduced, peak I_NaP_ is increased and I_NaR_ is diminished. As current traces in Figure 8B traces reveal, inversely adjusting I_NaR_ and I_NaP_ in this way does not affect recovery of fast inactivated channels into the open state, i.e., channels are still able to recover from fast inactivation into an open/conducting state on repolarization of the membrane.

**Figure 8.**
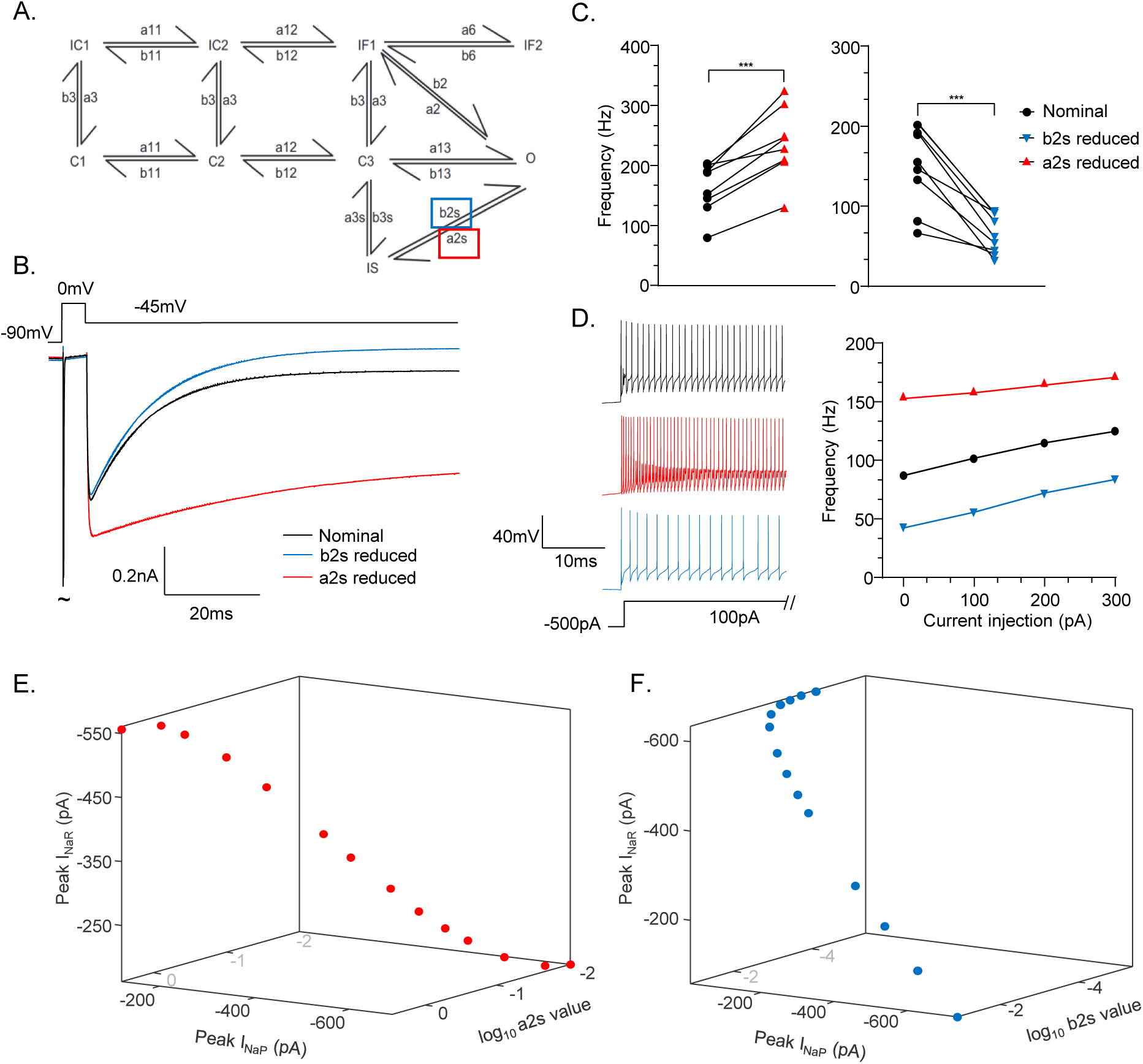
Peak I_NaP_ and I_NaR_ can be inversely adjusted by changing rate constants responsible for slow inactivation occupancy **A.** To develop models with inversely scaled I_NaP_ and I_NaR_, the state transition rate constants between the open (O) and slow inactivated (IS) states were individually altered. Decreasing the b2s (*blue box*) transition rate constant (in the nominal model) resulted in a conductance model with increased I_NaR_ and decreased I_NaP_. Decreasing the a2s (*red box*) transition rate constant resulted in a conductance model with decreased I_NaR_ and increased I_NaP_. These changes resulted in the simulated voltage-clamp traces that elicit the b2s reduced (*blue*) and a2s reduced (red) models, which are compared to the nominal model (*black*) in panel **B. C.** When compared to the nominal conductance model (black circles), spontaneous firing frequency of adult Purkinje neurons was significantly *increased* after addition of the a2s reduced model (*red triangles, left*), and significantly *decreased* after addition of the b2s reduced model (*blue triangles, right*) (paired Student’s t-test. 400 nS a2s reduced, P = 0.0004, n = 8; b2s reduced, P = 0.0029, n = 9;). **D.** Representative recordings of action potentials during addition of the nominal model (*black, top*), the a2s decreased (*red, middle*), and b2s decreased (*blue, bottom*) models (*left*) reveal scaling in spontaneous and evoked (see F-I plots, *right*) firing rates in a single Purkinje neuron. **E. F.** 3D plots were created in which the a2s (E.) or b2s (F.) state transition rate constants are decreased across 14 iterations and plotted as log10 against the resulting peak I_NaP_ and I_NaR_ values (measured at -45 mV).

Previous dynamic clamp experiments suggest peak I_NaR_ has no significant impact on the firing rate or action potential waveform properties of cerebellar Purkinje neurons. However, in adjusting rate constants that affect occupancy in the slow inactivated state, we inversely scale I_NaR_ and I_NaP_. These data indicate the presence of I_NaR_ may reflect the degree of Nav channel slow inactivation, which could regulate or fine-tune the population of Nav channels available to participate in I_NaP_. We hypothesized that adjusting slow inactivation occupancy, and increasing or decreasing I_NaP_, as we have done if Figure 8B, will result in increased or decreased repetitive firing rates. To test this hypothesis, we applied the models developed in Figure 8B in dynamic clamp experiments, using application of the nominal model in dynamic clamp as a baseline. Indeed, as panels 8C and 8D show, adjusting the transition of open channels into the IS state can be used to inversely scale I_NaR_ and I_NaP_, which results in significant (compared to the nominal model) increases or decreases in both spontaneous (Fig. 8C) and evoked (Fig. 8D) repetitive firing frequencies. Plotting a2s (Fig. 8E) or b2s (Fig 8F) rate constants against the resulting Nav conductance models’ peak I_NaP_ and I_NaR_ further reveal this relationship. By decreasing the a2s state transition rate, and thereby decreasing accumulation into the slow inactivated (IS) state, peak I_NaP_ is increased while peak I_NaR_ is decreased. Alternatively, if the b2s state transition rate is decreased, accumulation into the slow inactivated state is increased, resulting in increased peak I_NaR_ and reduced I_NaP_.

## Discussion

We used dynamic clamp to apply several Nav conductance models (10), which have varying resurgent (I_NaR_) and persistent (I_NaP_) Nav current properties, to test how these current components contribute to the firing of adult cerebellar Purkinje neurons. Results from these experiments indicate peak I_NaR_ does not directly contribute to the repetitive firing rates of Purkinje neurons. Instead, the amplitude of I_NaP_, particularly at subthreshold voltages, reliably scales firing frequency regardless of I_NaR_ amplitude. Modeling work revealed I_NaR_ and I_NaP_ can be inversely scaled by adjusting occupancy in the slow inactivated kinetic state, whereby slow inactivation removes Nav channels from contributing to I_NaP_.

### I_NaR_ and excitability

In our initial experiments, we constructed two Markov kinetic state models with decreased or increased I_NaR_ by adjusting the properties of two pairs of state transition rate constants of a previously developed Nav conductance model (Ransdell et al., 2022). These hand-tuned Nav models were individually added to Purkinje neurons via dynamic clamp. Surprisingly, after adding a model Nav conductance with an increased I_NaR_ component, there were no significant changes in firing frequency compared to the base (nominal) model (Figure 2). Additional dynamic clamp studies utilized Nav models which differ in the peak amplitude of I_NaR_ and I_NaP_, or that vary in the time constant of I_NaR_ decay. These studies revealed I_NaP_ is the critical Nav conductance parameter for tuning Purkinje neuron repetitive firing frequency, while peak I_NaR_ has very little effect on firing frequency or the action potential waveform. Of course, this provokes an important question— what role does I_NaR_ serve in regulating Purkinje neuron function if it has no effect on action potential firing? We propose I_NaR_, or more specifically the decay of I_NaR_ (which is rapid at subthreshold voltages), may reflect the removal of non-inactivating Nav channels into a slow inactivated kinetic state, and thus, I_NaR_ reflects a mechanism by which Purkinje neuron I_NaP_ is regulated. Alternative and/or additional potential contributions of I_NaR_ to Purkinje neuron excitability exist. Outside of the sustained repetitive simple spike firing characterized here, Purkinje neurons in the acute slice preparation have been shown to fire in trimodal patterns of activity which include trains of brief (<10 spikes) bursts of action potentials (M. D. Womack & Khodakhah, 2004; M. Womack & Khodakhah, 2002). Additionally, in response to excitatory drive from inferior olive climbing fibers (Eccles et al., 1966; Fujita, 1968), Purkinje neurons intermittently fire complex spikes, which involve an initial axonal spike followed by high frequency spikelets lasting several milliseconds (Stuart & Häusser, 1994). The sustained depolarization necessary for both Purkinje neuron bursting and complex spikes may rely on the contribution of I_NaR._

There is a long-standing hypothesis that I_NaR_ increases membrane excitability and scales neuronal repetitive firing rates since its discovery by Raman and Bean in 1997. *Scn8a* mutant mice (*Scn8a* null) have reduced I_NaR_ (by 90%) and reduced Purkinje neuron firing frequencies, which suggests I_NaR_ contributes to repetitive action potential generation (Raman et al., 1997). However, it was also noted that the targeted deletion of *Scn8a* results in reduced (by ∼70%) I_NaP_, which may have caused the attenuated repetitive firing rates (see Figures 4, 8). *Scn8a* null mice were also found to have hyperpolarized interspike voltages, which also might reflect attenuated subthreshold I_NaP_ (Raman et al., 1997).

The decay of I_NaR_ at subthreshold voltages is rapid (∼3 ms) (Raman & Bean, 1997; Zemel et al., 2021), suggesting the majority of depolarizing Nav current during interspike intervals, and the Nav current immediately preceding action potentials during repetitive firing, is a result of subthreshold I_NaP_. Subthreshold Nav current has previously been shown to play a critical role in setting the rate of interspike interval depolarization as well as the action potential threshold voltage in mouse Purkinje neurons as well as CA1 hippocampal neurons (Carter et al., 2012). The Markov conductance models developed here and dynamic clamp studies presented in Figure 3 reveal that even if the time constant of I_NaR_ decay is increased (nearly doubled at -65 mV) or decreased, compared to the nominal model (Figure 3B), the effects of I_NaR_ on repetitive firing are similar (Figure 3E), revealing I_NaR_ does not directly influence repetitive firing frequency even with much slower decay kinetics.

Targeted deletion of *Scn4b*, which encodes the Nav channel β4 (Navβ4) subunit, results in reduced firing rates, as well as reduced (by ∼50%) peak I_NaR_ in Purkinje neurons (Ransdell et al., 2017). Importantly, this result was consistent with an earlier study which found the deletion of *Scn4b* causes reduced I_NaR_ and evoked firing rates in rat striatal medium spiny neurons (Miyazaki et al., 2014). To test if the deficits in *Scn4b^-/-^* Purkinje neuron firing could be rescued, Ransdell et al. (2017) applied a modeled I_NaR_ conductance via dynamic clamp to these cells and found that firing rates of *Scn4b^-/-^* Purkinje neurons could be rescued to wild type levels (Ransdell et al., 2017). Importantly, however, the model Nav conductance utilized in Ransdell et al., (2017) included a persistent Nav component and the effect of targeted *Scn4b* deletion on I_NaP_ was not investigated. In RA projection neurons (RAPNs) of the adult zebra finch, Zemel and colleagues present a collection of data showing that RAPN maturation includes paired increases in firing frequency and peak I_NaR_, which corresponded with increased levels of Navβ4 expression (Zemel et al., 2021). In these studies, dynamic clamp was again used to test the role of I_NaR_ on RAPN firing frequencies, however, similar to Ransdell et al. (2017), the I_NaR_ model used included a persistent Nav current component.

Scaling I_NaR_ across conductance models did not drive consistent changes in action potential duration (extended data Table 2-1). Previous work has shown Purkinje neurons express large and fast activating voltage-gated potassium currents which quickly repolarize the membrane during the action potential downstroke (Raman & Bean, 1999). These large hyperpolarizing conductances likely shunt I_NaR_ depolarization and prevent any consistent effect of I_NaR_ on action potential duration. This point underscores the reasonable assumption that I_NaR_ contribution to membrane excitability will vary across neuronal cell types with unique firing and ion channel expression properties (Lewis & Raman, 2014).

As with any dynamic clamp investigation which focuses on intact neurons, complex architecture and distinct neuronal compartments result in testing the effects of a conductance on a non-isopotential membrane. This may cause erroneous voltage measures, and as a result, erroneous dynamic clamp current injections. In our studies, we patched the broad end of the somatic Purkinje neuron membrane toward the axonal projection. Previously, it was shown through focal TTX application to either the peri-somatic region of the cell or at the first node of Ranvier along the axon, that action potential initiation and spontaneous firing in Purkinje neurons relies on somatic and peri-somatic Nav channels (Khaliq & Raman, 2006), suggesting the voltage measurements and corresponding dynamic clamp current injections in our studies were sufficient in affecting the action potential initiation zone of Purkinje neurons. Across cells included in the study, passive membrane properties did not correlate with the effect of the nominal Nav conductance on repetitive firing (see extended data Figure 1-2), suggesting firing was not impacted by variations in Purkinje neuron morphology. However, adding modeled Nav conductance (via a current injecting electrode in dynamic clamp) at a single point in the somatic membrane will not perfectly recapitulate the contribution of expressed Nav channels along the axon initial segment, which drive repetitive action potentials in Purkinje neurons. The results from the experiments performed here, however, were consistent in the indication that subthreshold I_NaP_ is the critical Nav current component for scaling repetitive firing rates across cells.

### Persistent current and its relationship with resurgent current

Since the discovery of the persistent sodium current in Purkinje neurons (Llinás & Sugimori, 1980), I_NaP_ has been implicated in promoting repetitive and complex firing in several neuronal cell types (Carter et al., 2012; Han et al., 2015; Lee & Heckman, 2001; Taddese & Bean, 2002; Yamanishi et al., 2018). Experiments here demonstrate subthreshold I_NaP_ is critical in scaling Purkinje neuron firing rate (Figure 6D), but also that I_NaR_ may reflect an important mechanism by which Nav channels are removed from contributing to I_NaP_. Increased levels of I_NaP_ and I_NaR_ have often been identified as a result of gain-of-function mutations in Nav channel α subunits which are associated with human diseases such as paroxysmal extreme pain disorder (PEPD), long QT syndrome, and epilepsies (Eberhardt et al., 2014; Hargus et al., 2013; Jarecki et al., 2010; Pan & Cummins, 2020). In addition to the increased levels of I_NaR_ and I_NaP_, several of the cells in these studies were found to have increased action potential firing (Estacion et al., 2008; Hargus et al., 2013; Pan & Cummins, 2020). Mutations in Nav channel pore-forming α subunits, which drive singular or dual changes in I_NaR_ and/or I_NaP_, provide important clues regarding how changes in Nav channel activation, fast inactivation, and slow inactivation processes affect these current components, and additionally, how alterations in I_NaR_ and/or I_NaP_ affect membrane excitability. For instance, PEPD-associated mutations in Nav1.7 typically impact the voltage-dependence of fast inactivation resulting in enhanced I_NaP_ (Dib-Hajj et al., 2013; Eberhardt et al., 2014). It will be interesting to examine how mutations in Nav channel pore-forming or accessory subunits affect I_NaR_ and I_NaP_ along with Nav channel slow inactivation processes in native neuronal cell types.

### Slow inactivation and adjusting the ratio of I_NaR_ and I_NaP_

Nav channel slow inactivation is distinct from conventional fast inactivation and occurs during prolonged periods of Nav channel activation, generally tens to hundreds of milliseconds (Aman & Raman, 2007; Silva, 2014). The Markov models (Ransdell et al., 2022) used here have parallel inactivation pathways, a fast inactivation pathway and slow inactivation pathway, used to accurately reproduce the fast inactivation kinetics of I_NaT_ and slower decay of I_NaR_ at steady-state voltages (see Figures 1A, B). This model topography (depicted in Figure 1B) is unique in that it does not rely on an ‘open-channel block’ kinetic state, which has previously been used to model and explain the gating mechanism of I_NaR_ (Raman & Bean, 2001). In the open-channel block hypothesis, I_NaR_ generation is thought to involve an extrinsic blocking particle that, on membrane depolarization, competes directly with the intrinsic fast inactivation gate of Nav channel α subunits. During membrane repolarization, the increased driving force on extracellular sodium is thought to knock this extrinsic blocking particle out of the channel pore allowing ‘resurgent’ sodium influx (I_NaR_) and an eventual accumulation of Nav channels into the conventional fast inactivated state (reflected as I_NaR_ decay) (Aman & Raman, 2010; Lewis & Raman, 2014; Raman & Bean, 2001). However, efforts to abolish I_NaR_ through the targeted deletion of potential extrinsic Nav channel blocking particles have not been successful (Miyazaki et al., 2014; Ransdell et al., 2017; White et al., 2019). Additionally, experimental analysis of how extracellular sodium concentration, as well as the voltage and duration of the initial membrane depolarization affect I_NaR_ have revealed inconsistencies with the open-channel block hypothesis (Ransdell et al., 2022).

The Markov models used here preserve I_NaR_ gating in the absence of an open-channel block kinetic state. On membrane repolarization, simulated channels in the nominal model recover from the conventional fast inactivated state into an open state, producing I_NaR_ and I_NaP_. The decay of I_NaR_ during sustained depolarization is driven by an accumulation of open and fast inactivated Nav channels into an absorbing slow inactivated kinetic state. Under this topography, channels failing to undergo conventional fast inactivation, a process which is most probable at around -45 mV, is responsible for both I_NaR_ and I_NaP_. Adjusting rate constants that determine entry into and out of the slow inactivated kinetic state readily tunes the portion of channels contributing to I_NaR_ and I_NaP_, which determines the model’s effect on Purkinje neuron firing (Figure 8). From these results, it seems Nav channels can be regulated in at least two domains in order to regulate repetitive firing frequency: 1. the proportion of channels that are activated and fail to undergo fast inactivation (I_NaR_ + I_NaP_) and 2. the proportion of these non-inactivating channels which undergo slow inactivation (I_NaR_). These domains of Nav channel regulation would regulate the level of subthreshold I_NaP_ during interspike intervals, which, as our dynamic clamp studies show, scales Purkinje neuron firing rates.

Slow inactivation of Nav currents has previously been shown to be essential in the regulation of pacemaking activity in STN neurons (Do & Bean, 2003) and interspike voltage changes (Carter et al., 2012). Our results further suggest the magnitude of I_NaP_ may be regulated by Nav channel slow inactivation in Purkinje neurons. Enhancing cAMP-dependent protein kinase and protein kinase C activity has been shown to rapidly enhance Nav channel slow inactivation in heterologously expressed Nav1.2 channels (Chen et al., 2006), suggesting post-translational regulation of Nav channels can quickly modify this inactivation pathway. Reduced extracellular Na^+^ concentrations have also been found to increase entry into the slow inactivated kinetic state (Afshari et al., 2004; Aman & Raman, 2007; Townsend & Horn, 1997). Moving forward, it will be interesting to test how the regulation of slow inactivation processes in native neuronal cell types affect Nav current components, particularly I_NaR_ and I_NaP_, and to determine if post-translational regulation of Nav channel slow inactivation is capable/necessary for the fine-tuning of I_NaP_ as well as firing rates in spontaneously active cells.

## Supporting information

Extended Data Figures Brown et al.

## Acknowledgments

This work was supported by startup funds from the Miami University College of Arts and Sciences and by the NINDS at the National Institutes of Health: Award 1R15NS125560

## Author Contributions

SPB and JLR conceived and planned the studies; SPB and RJL performed dynamic clamp experiments; SPB and JDM developed Nav conductance models for dynamic clamp; SPB and JLR wrote the manuscript; SPB and JLR revised the manuscript.

## Extended Data Figure Legends

**Extended Data Figure 1-1.** Frequency-current relationship for Purkinje neurons with no dynamic clamp (open circles; n = 24) and after application of all Markov models (nominal, black circles, n = 24; τdecay decreased, green squares, n = 31; τdecay increased, blue triangles, n = 33; INaR reduced, orange triangles, n = 34; INaP reduced, red diamonds, n = 34; INaR reduced (hand-tuned), open grey circles, n = 10; INaR increased (hand-tuned), open squares, n = 9).

**Extended Data Figure 1-2.** Correlation between cell properties and the percent increase in firing. Capacitance (left) and resistance (right) is plotted against the percent increase in firing frequency after the nominal dynamic clamp model was applied. Each correlation is fitted with a simple linear regression and the correlation coefficients for capacitance and resistance are 0.024 (n = 28) and 0.012 (n = 27), respectively.

**Extended Data Table 2-1.** Correlations between the change in action potential threshold, duration, peak, amplitude, afterhyperpolarization, and max dv/dt before and after addition of dynamic clamp models plotted against the peak simulated values of INaP at -65 mV (left column), INaR at -45 mV (middle column), and INaT at -30 mV (right column) for each model. Correlation coefficients (r2) are generated using a simple linear regression. The asterisk denotes the r2 value of a linear regression fit of the data that includes the ‘INaR increased (hand-tuned)’ model, while the r2 value above is after removal of the ‘INaR increased (hand-tuned)’ data point.

## Notes

https://github.com/morenomdphd/Resurgent_INa

### Competing Interest Statement

The authors have declared no competing interest.

### Summary of Updates

This version of the manuscript has been revised based on comments from reviewers at the Journal of Neuroscience.

## References

1. Afshari, F. S., Ptak, K., Khaliq, Z. M., Grieco, T. M., Slater, N. T., McCrimmon, D. R., & Raman, I. M. (2004). Resurgent Na Currents in Four Classes of Neurons of the Cerebellum. Journal of Neurophysiology, 92(5), 2831–2843. 10.1152/jn.00261.2004

2. Aman, T. K., & Raman, I. M. (2007). Subunit dependence of Na channel slow inactivation and open channel block in cerebellar neurons. Biophysical Journal, 92(6), 1938–1951. 10.1529/biophysj.106.093500

3. Aman, T. K., & Raman, I. M. (2010). Inwardly permeating Na ions generate the voltage dependence of resurgent Na current in cerebellar Purkinje neurons. The Journal of Neuroscience: The Official Journal of the Society for Neuroscience, 30(16), 5629–5634. 10.1523/JNEUROSCI.0376-10.2010

4. Carter, B. C., Giessel, A. J., Sabatini, B. L., & Bean, B. P. (2012). Transient sodium current at subthreshold voltages: Activation by EPSP waveforms. Neuron, 75(6), 1081–1093. 10.1016/j.neuron.2012.08.033

5. Chen, Y., Yu, F. H., Surmeier, D. J., Scheuer, T., & Catterall, W. A. (2006). Neuromodulation of Na+ channel slow inactivation via cAMP-dependent protein kinase and protein kinase C. Neuron, 49(3), 409–420. 10.1016/j.neuron.2006.01.009

6. Dib-Hajj, S. D., Yang, Y., Black, J. A., & Waxman, S. G. (2013). The Na(V)1.7 sodium channel: From molecule to man. Nature Reviews. Neuroscience, 14(1), 49–62. 10.1038/nrn3404

7. Do, M. T. H., & Bean, B. P. (2003). Subthreshold sodium currents and pacemaking of subthalamic neurons: Modulation by slow inactivation. Neuron, 39(1), 109–120. 10.1016/s0896-6273(03)00360-x

8. Eberhardt, M., Nakajima, J., Klinger, A. B., Neacsu, C., Hühne, K., O’Reilly, A. O., Kist, A. M., Lampe, A. K., Fischer, K., Gibson, J., Nau, C., Winterpacht, A., & Lampert, A. (2014). Inherited pain: Sodium channel Nav1.7 A1632T mutation causes erythromelalgia due to a shift of fast inactivation. The Journal of Biological Chemistry, 289(4), 1971–1980. 10.1074/jbc.M113.502211

9. Eccles, J. C., Llinás, R., & Sasaki, K. (1966). The excitatory synaptic action of climbing fibres on the Purkinje cells of the cerebellum. The Journal of Physiology, 182(2), 268–296. 10.1113/jphysiol.1966.sp007824

10. Estacion, M., Dib-Hajj, S. D., Benke, P. J., Te Morsche, R. H. M., Eastman, E. M., Macala, L. J., Drenth, J. P. H., & Waxman, S. G. (2008). NaV1.7 gain-of-function mutations as a continuum: A1632E displays physiological changes associated with erythromelalgia and paroxysmal extreme pain disorder mutations and produces symptoms of both disorders. The Journal of Neuroscience: The Official Journal of the Society for Neuroscience, 28(43), 11079–11088. 10.1523/JNEUROSCI.3443-08.2008

11. Fujita, Y. (1968). Activity of dendrites of single Purkinje cells and its relationship to so-called inactivation response in rabbit cerebellum. Journal of Neurophysiology, 31(2), 131–141. 10.1152/jn.1968.31.2.131

12. Han, C., Estacion, M., Huang, J., Vasylyev, D., Zhao, P., Dib-Hajj, S. D., & Waxman, S. G. (2015). Human Nav1.8: Enhanced persistent and ramp currents contribute to distinct firing properties of human DRG neurons. Journal of Neurophysiology, 113(9), 3172– 3185. 10.1152/jn.00113.2015

13. Hargus, N. J., Nigam, A., Bertram, E. H., & Patel, M. K. (2013). Evidence for a role of Nav1.6 in facilitating increases in neuronal hyperexcitability during epileptogenesis. Journal of Neurophysiology, 110(5), 1144–1157. 10.1152/jn.00383.2013

14. Häusser, M., & Clark, B. A. (1997). Tonic synaptic inhibition modulates neuronal output pattern and spatiotemporal synaptic integration. Neuron, 19(3), 665–678. 10.1016/s0896-6273(00)80379-7

15. Ito, M. (1984). The Cerebellum and Neural Control (1st ed.). Raven Press.

16. Ito, M., Yoshida, M., & Obata, K. (1964). Monosynaptic inhibition of the intracerebellar nuclei induced from the cerebellar cortex. Experientia, 20(10), 575–576. 10.1007/BF02150304

17. Jarecki, B. W., Piekarz, A. D., Jackson, J. O., & Cummins, T. R. (2010). Human voltage-gated sodium channel mutations that cause inherited neuronal and muscle channelopathies increase resurgent sodium currents. The Journal of Clinical Investigation, 120(1), 369–378. 10.1172/JCI40801

18. Khaliq, Z. M., Gouwens, N. W., & Raman, I. M. (2003). The contribution of resurgent sodium current to high-frequency firing in Purkinje neurons: An experimental and modeling study. The Journal of Neuroscience: The Official Journal of the Society for Neuroscience, 23(12), 4899–4912. 10.1523/JNEUROSCI.23-12-04899.2003

19. Khaliq, Z. M., & Raman, I. M. (2006). Relative Contributions of Axonal and Somatic Na Channels to Action Potential Initiation in Cerebellar Purkinje Neurons. The Journal of Neuroscience, 26(7), 1935–1944. 10.1523/JNEUROSCI.4664-05.2006

20. Lagarias, J. C., Reeds, J. A., Wright, M. H., & Wright, P. E. (1998). Convergence Properties of the Nelder—Mead Simplex Method in Low Dimensions. SIAM Journal on Optimization, 9(1), 112–147. 10.1137/S1052623496303470

21. Lee, R. H., & Heckman, C. J. (2001). Essential role of a fast persistent inward current in action potential initiation and control of rhythmic firing. Journal of Neurophysiology, 85(1), 472–475. 10.1152/jn.2001.85.1.472

22. Lewis, A. H., & Raman, I. M. (2014). Resurgent current of voltage-gated Na+ channels. The Journal of Physiology, 592(Pt 22), 4825–4838. 10.1113/jphysiol.2014.277582

23. Llinás, R., & Sugimori, M. (1980). Electrophysiological properties of in vitro Purkinje cell somata in mammalian cerebellar slices. The Journal of Physiology, 305, 171–195. 10.1113/jphysiol.1980.sp013357

24. Miyazaki, H., Oyama, F., Inoue, R., Aosaki, T., Abe, T., Kiyonari, H., Kino, Y., Kurosawa, M., Shimizu, J., Ogiwara, I., Yamakawa, K., Koshimizu, Y., Fujiyama, F., Kaneko, T., Shimizu, H., Nagatomo, K., Yamada, K., Shimogori, T., Hattori, N., … Nukina, N. (2014). Singular localization of sodium channel β4 subunit in unmyelinated fibres and its role in the striatum. Nature Communications, 5, 5525. 10.1038/ncomms6525

25. Moreno, J. D., Lewis, T. J., & Clancy, C. E. (2016). Parameterization for In-Silico Modeling of Ion Channel Interactions with Drugs. PLOS ONE, 11(3), e0150761. 10.1371/journal.pone.0150761

26. Nelder, J. A., & Mead, R. (1965). A Simplex Method for Function Minimization. The Computer Journal, 7(4), 308–313. 10.1093/comjnl/7.4.308

27. Obata, K., Ito, M., Ochi, R., & Sato, N. (1967). Pharmacological properties of the postsynaptic inhibition by Purkinje cell axons and the action of γ-aminobutyric acid on Deiters neurones. Experimental Brain Research, 4(1), 43–57. 10.1007/BF00235216

28. Obata, K., Takeda, K., & Shtnozaki, H. (1970). Further study on pharmacological properties of the cerebellar-induced inhibition of Deiters neurones. Experimental Brain Research, 11(4), 327–342. 10.1007/BF00237907

29. Pan, Y., & Cummins, T. R. (2020). Distinct functional alterations in SCN8A epilepsy mutant channels. The Journal of Physiology, 598(2), 381–401. 10.1113/JP278952

30. Quattrocolo, G., Dunville, K., & Nigro, M. J. (2021). Resurgent Sodium Current in Neurons of the Cerebral Cortex. Frontiers in Cellular Neuroscience, 15, 760610. 10.3389/fncel.2021.760610

31. Raman, I. M. (2023). The Hodgkin-Huxley-Katz Prize Lecture: A Markov model with permeation-dependent gating that accounts for resurgent current of voltage-gated Na channels. The Journal of Physiology. 10.1113/JP285166

32. Raman, I. M., & Bean, B. P. (1997). Resurgent sodium current and action potential formation in dissociated cerebellar Purkinje neurons. The Journal of Neuroscience: The Official Journal of the Society for Neuroscience, 17(12), 4517–4526. 10.1523/JNEUROSCI.17-12-04517.1997

33. Raman, I. M., & Bean, B. P. (1999). Ionic currents underlying spontaneous action potentials in isolated cerebellar Purkinje neurons. The Journal of Neuroscience: The Official Journal of the Society for Neuroscience, 19(5), 1663–1674. 10.1523/JNEUROSCI.19-05-01663.1999

34. Raman, I. M., & Bean, B. P. (2001). Inactivation and recovery of sodium currents in cerebellar Purkinje neurons: Evidence for two mechanisms. Biophysical Journal, 80(2), 729–737. 10.1016/S0006-3495(01)76052-3

35. Raman, I. M., Sprunger, L. K., Meisler, M. H., & Bean, B. P. (1997). Altered subthreshold sodium currents and disrupted firing patterns in Purkinje neurons of Scn8a mutant mice. Neuron, 19(4), 881–891. 10.1016/s0896-6273(00)80969-1

36. Ransdell, J. L., Dranoff, E., Lau, B., Lo, W.-L., Donermeyer, D. L., Allen, P. M., & Nerbonne, J. M. (2017). Loss of Navβ4-Mediated Regulation of Sodium Currents in Adult Purkinje Neurons Disrupts Firing and Impairs Motor Coordination and Balance. Cell Reports, 19(3), 532–544. 10.1016/j.celrep.2017.03.068

37. Ransdell, J. L., Moreno, J. D., Bhagavan, D., Silva, J. R., & Nerbonne, J. M. (2022). Intrinsic mechanisms in the gating of resurgent Na+ currents. eLife, 11, e70173. 10.7554/eLife.70173

38. Ransdell, J. L., & Nerbonne, J. M. (2018). Voltage-gated sodium currents in cerebellar Purkinje neurons: Functional and molecular diversity. Cellular and Molecular Life Sciences: CMLS, 75(19), 3495–3505. 10.1007/s00018-018-2868-y

39. Silva, J. (2014). Slow inactivation of Na(+) channels. Handbook of Experimental Pharmacology, 221, 33–49. 10.1007/978-3-642-41588-3_3

40. Solinas, S., Forti, L., Cesana, E., Mapelli, J., De Schutter, E., & D’Angelo, E. (2007). Computational reconstruction of pacemaking and intrinsic electroresponsiveness in cerebellar Golgi cells. Frontiers in Cellular Neuroscience, 1, 2. 10.3389/neuro.03.002.2007

41. Stuart, G., & Häusser, M. (1994). Initiation and spread of sodium action potentials in cerebellar Purkinje cells. Neuron, 13(3), 703–712. 10.1016/0896-6273(94)90037-x

42. Taddese, A., & Bean, B. P. (2002). Subthreshold Sodium Current from Rapidly Inactivating Sodium Channels Drives Spontaneous Firing of Tuberomammillary Neurons. Neuron, 33(4), 587–600. 10.1016/S0896-6273(02)00574-3

43. Thach, W. T. (1968). Discharge of Purkinje and cerebellar nuclear neurons during rapidly alternating arm movements in the monkey. Journal of Neurophysiology, 31(5), 785–797. 10.1152/jn.1968.31.5.785

44. Townsend, C., & Horn, R. (1997). Effect of alkali metal cations on slow inactivation of cardiac Na+ channels. The Journal of General Physiology, 110(1), 23–33. 10.1085/jgp.110.1.23

45. White, H. V., Brown, S. T., Bozza, T. C., & Raman, I. M. (2019). Effects of FGF14 and NaVβ4 deletion on transient and resurgent Na current in cerebellar Purkinje neurons. Journal of General Physiology, 151(11), 1300–1318. 10.1085/jgp.201912390

46. Womack, M. D., & Khodakhah, K. (2004). Dendritic Control of Spontaneous Bursting in Cerebellar Purkinje Cells. Journal of Neuroscience, 24(14), 3511–3521. 10.1523/JNEUROSCI.0290-04.2004

47. Womack, M., & Khodakhah, K. (2002). Active Contribution of Dendrites to the Tonic and Trimodal Patterns of Activity in Cerebellar Purkinje Neurons. Journal of Neuroscience, 22(24), 10603–10612. 10.1523/JNEUROSCI.22-24-10603.2002

48. Yamanishi, T., Koizumi, H., Navarro, M. A., Milescu, L. S., & Smith, J. C. (2018). Kinetic properties of persistent Na+ current orchestrate oscillatory bursting in respiratory neurons. The Journal of General Physiology, 150(11), 1523–1540. 10.1085/jgp.201812100

49. Zemel, B. M., Nevue, A. A., Dagostin, A., Lovell, P. V., Mello, C. V., & von Gersdorff, H. (2021). Resurgent Na+ currents promote ultrafast spiking in projection neurons that drive fine motor control. Nature Communications, 12(1), 6762. 10.1038/s41467-021-26521-3

