## Extended Data Figures Brown et al. for "A Reinterpretation of the Relationship Between Persistent and Resurgent Sodium Currents"

### Extended Data Figure 1-1. Brown et al.

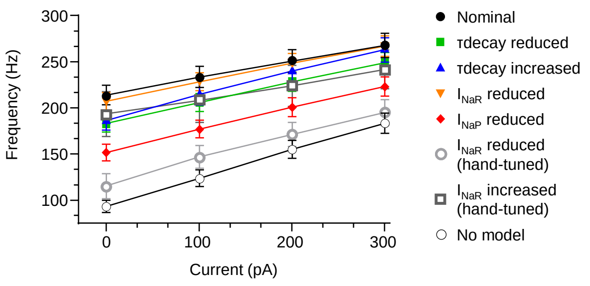


**Figure 1-1.** Frequency-current relationship for Purkinje neurons with no dynamic clamp (open circles; n = 24) and after application of all Markov models (nominal, black circles, n = 24; τdecay decreased, green squares, n = 31; τdecay increased, blue triangles, n = 33; INaR reduced, orange triangles, n = 34; INaP reduced, red diamonds, n = 34; INaR reduced (hand-tuned), open grey circles, n = 10; INaR increased (hand-tuned), open squares, n = 9).

### Extended Data Figure 1-2. Brown et al.

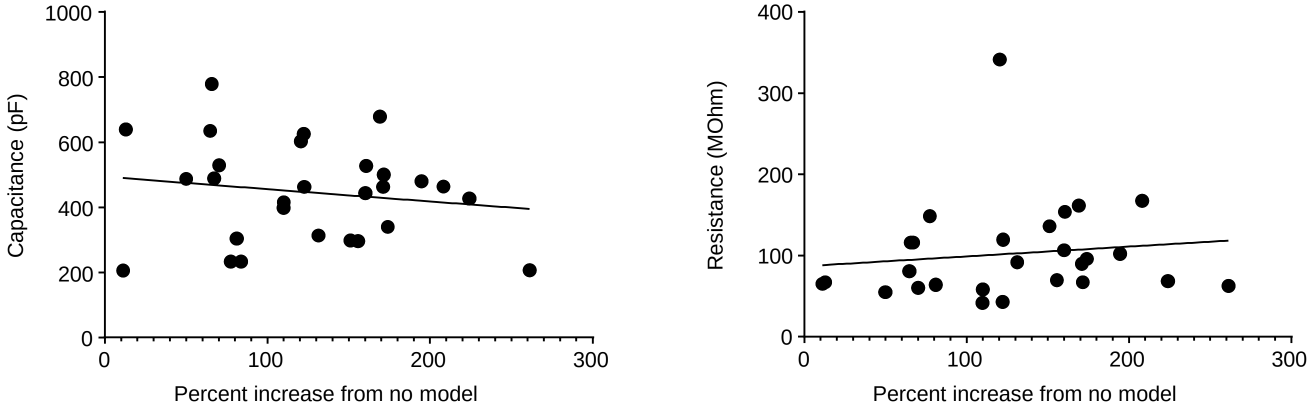


**Figure 1-2.** Correlation between cell properties and the percent increase in firing. Capacitance (left) and resistance (right) is plotted against the percent increase in firing frequency after the nominal dynamic clamp model was applied. Each correlation is fitted with a simple linear regression and the correlation coefficients for capacitance and resistance are 0.024 (n = 28) and 0.012 (n = 27), respectively.

### Extended Data Table 1-1. Brown et al.

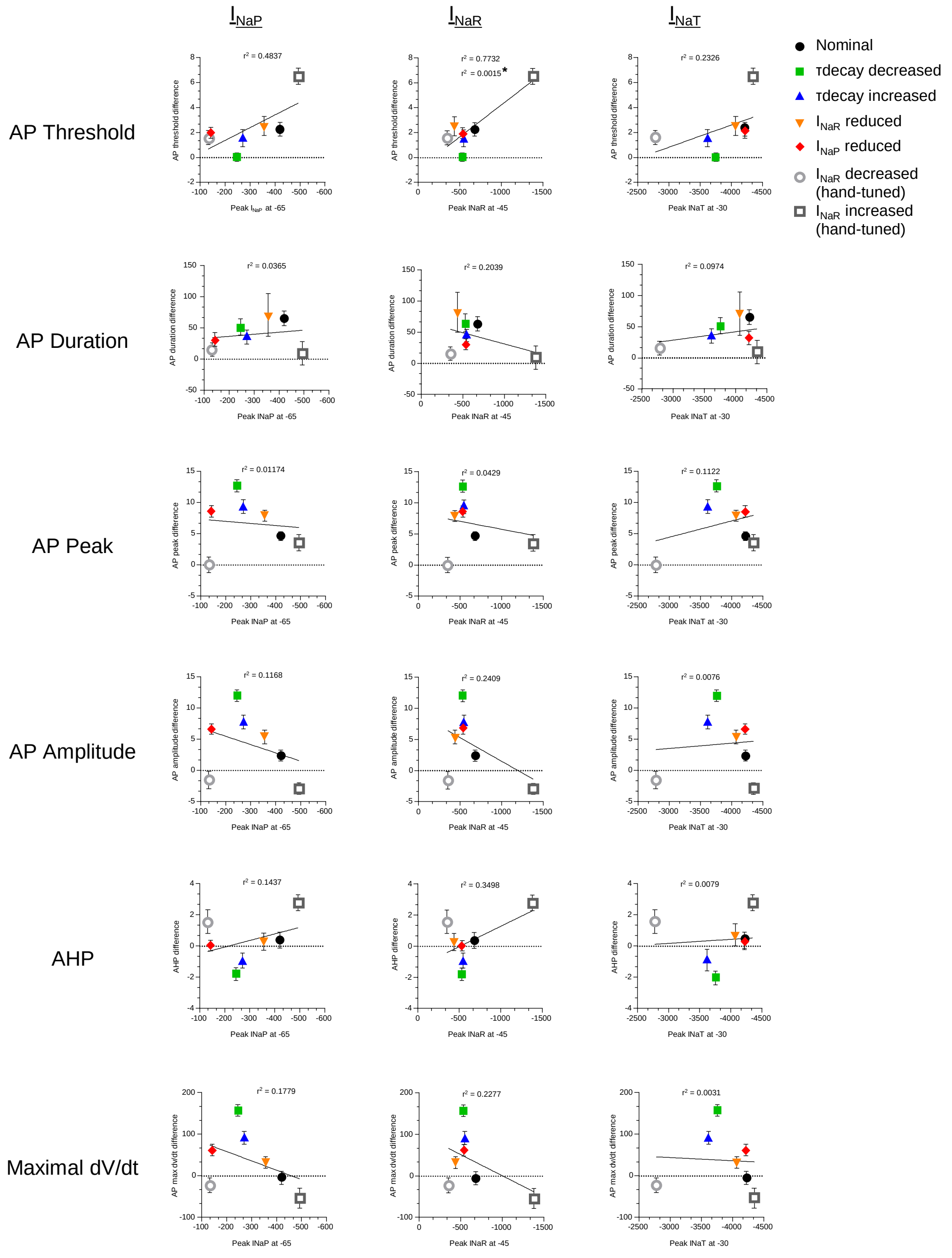


**Extended Data Table 1-1.** Correlation between the difference of action potential threshold, duration, peak, amplitude, afterhyperpolarization, and max dv/dt before and after addition of dynamic clamp models and the peak simulated values of INaP at -65 mV (left column), INaR at -45 mV (middle column), and INaT at -30 mV (right column) for each model. Correlation coefficients (r2) are generated by a simple linear regression. The asterisk denotes the r2 value of a linear regression fit of the data that includes the ‘INaR increased (hand-tuned)’ model, while the r2 value above is with that data point removed.
